# Cellular and network mechanisms may generate sparse coding of sequential object encounters in hippocampal-like circuits

**DOI:** 10.1101/571414

**Authors:** Anh-Tuan Trinh, Stephen E. Clarke, Erik Harvey-Girard, Leonard Maler

## Abstract

In mammals, the localization of distinct landmarks is performed by hippocampal neurons that sparsely encode an animal’s location relative to surrounding objects. Similarly, the dorsal lateral pallium (DL) is essential for spatial learning in teleost fish. The DL of weakly electric gymnotiform fish receives sensory inputs from the preglomerular nucleus (PG), which has been hypothesized to encode the temporal sequence of electrosensory or visual landmark/food encounters. Here, we show that DL neurons have a hyperpolarized resting membrane potential combined with a high and dynamic spike threshold that increases following each spike. Current-evoked spikes in DL cells are followed by a strong small-conductance calcium-activated potassium channel (SK) mediated after-hyperpolarizing potential (AHP). Together, these properties prevent high frequency and continuous spiking. The resulting sparseness of discharge and dynamic threshold suggest that DL neurons meet theoretical requirements for generating spatial memory engrams by decoding the landmark/food encounter sequences encoded by PG neurons.

## Introduction

The mammalian hippocampus is required for the storage and recall of various types of memory, including spatial memory that presumably guides path integration and landmark based navigation (Barry and Burgess, 2014; Hartley et al., 2014). In the conventional view, sparse discharge of dentate gyrus (DG) granule cells and CA1/CA3 pyramidal cells can encode a rodent’s location (*i.e.*, place field) with respect to visually identified landmarks (Barry and Burgess, 2014; Hartley et al., 2014). An emerging alternate view of hippocampal function emphasizes its role in the encoding of temporal sequences within or across periods of locomotion (Pastalkova et al., 2008; MacDonald et al., 2011; Kraus et al., 2013; Eichenbaum, 2014; Modi et al., 2014; Ranganath and Hsieh, 2016). For example, hippocampal neurons may discharge at specific times after the initiation of running on a treadmill and effectively tile an entire running episode (Kraus et al., 2013). The encoding of time and location appears to be closely connected with the responses of a subset of neurons to time spent and distance travelled (Kraus et al., 2013; Deuker et al., 2016; Eichenbaum, 2017).

Visuospatial memory is also important for teleost fish (Rodriguez et al., 2002) and they can learn to finely discriminate between visual inputs (Schluessel and Bleckmann, 2005; Siebeck et al., 2009; Rischawy and Schuster, 2013; Newport et al., 2016). Unlike mammals, fish do not have an obvious cortex or hippocampus; instead, their dorsal telencephalon (pallium) is divided into non-layered cell groups that have specific connectivity and function (Rodríguez et al., 2002; Northcutt, 2008; Giassi et al., 2012c; Giassi et al., 2012b). Visual input to the pallium primarily arrives from the optic tectum and reaches the dorsolateral pallium (DL) through the thalamus-like preglomerular nucleus (PG, Yamamoto and Ito, 2008; Giassi et al., 2012c; Wallach et al., 2018). Lesion studies have shown that DL is essential for visual (landmark) based spatial learning and memory (Rodriguez et al., 2002).

Comparisons of teleost pallium and mammalian hippocampus and cortex have been controversial, and similarity between DL and either hippocampus or cortex have been stressed. Based on its location (Yamamoto et al., 2007; Mueller and Wullimann, 2009), extrinsic connections (Elliott et al., 2017) and molecular markers (Harvey-Girard et al., 2012; Ganz et al., 2014), it has been proposed that DL is homologous to the hippocampus (in particular to DG, Elliott et al., 2017). However, unlike the major recipients of sensory information in the hippocampal formation (via cortex, *i.e.*, DG, CA1), DL neurons have strong local recurrent connectivity (Trinh et al., 2016). The extrinsic and intrinsic connectivity of DL also suggests a strong resemblance to the mammalian cortex (Yamamoto et al., 2007; Giassi et al., 2012c; Trinh et al., 2016; Elliott et al., 2017). In either case, DL neurons are morphologically very different from both DG granule cells and the pyramidal cells of the hippocampus and cortex: they lack long apical dendrites and, instead, have short isotropic dendrites (Giassi et al., 2012b).

A teleost subgroup, the weakly electric gymnotiform fish, have poor vision and rely heavily on their topographically organized electrosense for navigation (Krahe et al., 2008; Fotowat et al., 2018) and electrolocation; (Clarke et al., 2015). They can use their electrosensory system to finely discriminate temporal (Harvey-Girard et al., 2010) and spatial (Graff et al., 2004; Dangelmayer et al., 2016) patterns and use electrosensory-identified landmarks to learn the spatial location of food in the dark (Jun et al., 2016). Electrosensory input is first processed by a topographically mapped 2-D cutaneous electroreceptor array found in the hindbrain electrosensory lobe (ELL) and, via a midbrain relay, then mapped onto the tectum (Krahe and Maler, 2014); electrosensory and visual input are even combined in a subset of tectal neurons (Bastian, 1982). Electrosensory and visual tectal cells then project to PG and their PG target then projects exclusively to DL (Giassi et al., 2012c).

A recent paper has demonstrated similar adaptive coding properties of PG cells for visual and electrosensory motion stimuli (Wallach et al., 2018). Two recent studies have shown that DL cells process this input. In a gymnotiform fish, neurons within a major target of DL (dorsal pallium, DD) have been shown to discharge to the electrosensory signals generated when the fish moves near ‘landmarks’ (Fotowat et al., 2018). In goldfish, Vinepinsky et al. (2018) have described DL cells responsive to boundaries (visual input) as well as speed and direction of self-motion. Given this data, and the similar anatomical and functional organization of visual and electrosensory motion pathways, we hypothesize that electrosensory motion signals are processed in DL to generate spatial memories that are also stored in DL (Fig. 1).

**Figure 1.**
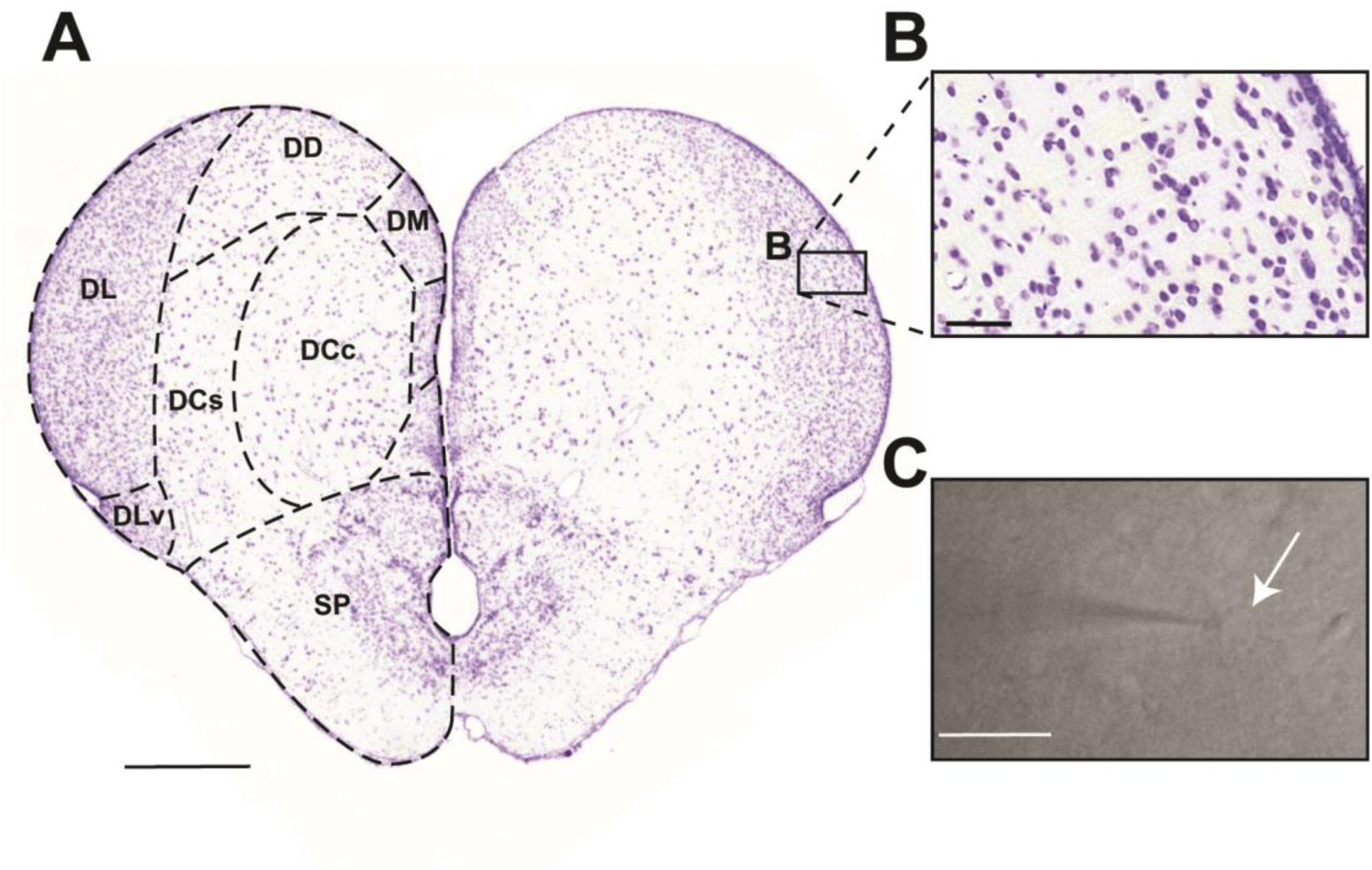
Anatomy of the *Apteronotus leptorhynchus* telencephalon. **A**. A transverse section through the *Apteronotus* telencephalon indicating the major subdivisions of pallium and subpallium; this section was obtained from a standard series of cresyl violet stained sections (Elliot et al, 2017). Midbrain sensory inputs entering the pallium from PG terminates in the dorsolateral pallium (DL). These inputs are processed within the DL recurrent network (Trinh et al., 2016). DL projects to the core dorsalcentral pallium (DCc) which, in turn, projects to midbrain sensory regions. DLv is located ventral to DL and distinguished by its olfactory bulb input. The dorsal-dorsal pallium (DD) has reciprocal connections with DL (Elliott et al., 2017). Scale bar: 500 μm. **B**. A higher magnification of the cells in DL illustrates an apparent random distribution and its highly organized intrinsic laminar and columnar circuitry is not evident (Trinh et al., 2016). The neurons in DL have homogenous morphology and are roughly 10 μm in diameter (Giassi et al., 2012b). Scale bar: 50 μm. **C**. An infrared image of a DL neuron undergoing a whole-cell patch recording. The shadow to the left illustrates the patch pipette, while the white arrow highlights the patched cell. Scale bar: 20 μm. **Abbreviations**
DCc: dorsocentral pallium, core
DCs: dorsocentral pallium, shell
DD: dorsodorsal pallium
DL: dorsolateral pallium
DLv: dorsolateral pallium, ventral subdivision
DM: dorsomedial pallium
SP: subpallium

A subset of electrosensory motion PG neurons have been identified that can encode, via their firing rate, the time interval between object encounters (Wallach et al., 2018). This led Wallach et al. (2018) to hypothesize that the output of these ‘time stamp’ neurons is used to estimate the distance between the objects encountered by the fish, thereby supporting the observed electrosense-dependent spatial learning (Jun et al., 2016). We hypothesize that the transformation of the PG derived ‘time stamp’ electrosensory information to a spatial map takes place in DL (Giassi et al., 2012c). Here, we studied the biophysical properties of DL neurons *in vitro* to determine whether their intrinsic properties are compatible with their putative role in converting temporal input from PG (*i.e.*, time between object encounters) to a spatial map (Wallach et al., 2018).

## Results

We performed whole cell patch recordings from *Apteronotus* DL neurons in acute slices from the rostral- to mid-telencephalon (Fig. 1A). Cells within DL, imaged under infrared illumination with DIC optics, had a shape and size consistent with those identified in Nissl stained sections (Fig. 1B, C). Although we cannot differentiate between excitatory and inhibitory cells, we assume that the neurons whose biophysical properties we characterize are almost certainly those of excitatory (glutamatergic) DL neurons since they vastly predominate over the rare inhibitory (GABAergic) cells (Giassi et al., 2012b). We also recorded neurons from the dorsal portion of *Carassius auratus* (goldfish) DL, while avoiding the ventral DL as it receives olfactory bulb input (Northcutt, 2006). The physiology of neurons recorded in the goldfish DL was not distinguishable from those of *Apteronotus* (see below).

### ‘Noisy’ versus ‘quiet’ cells

After attaining the whole-cell patch configuration, we first examined the resting membrane potential (RMP, no holding current), and observed two distinct electrophysiological profiles. The majority of the DL cells (29/35 cells in *Apteronotus* and 7/11 cells in goldfish) were ‘quiet’, that is, they had minimal spontaneous membrane fluctuations, as shown by the example recording traces from three different *Apteronotus* DL cells with different RMPs (Fig 2A). A smaller number of DL neurons were ‘noisy,’ showing considerable spontaneous membrane fluctuations over approximately the same range of RMPs as the ‘quiet’ cells (Fig 2B). A histogram estimating the distribution of RMP variance (Fig. 2C) suggests that, in both *Apteronotus* and goldfish DL, there were distinct populations of quiet (variance < 0.5 mV^2^) and noisy cells (variance > 0.5 mV^2^).

**Figure 2.**
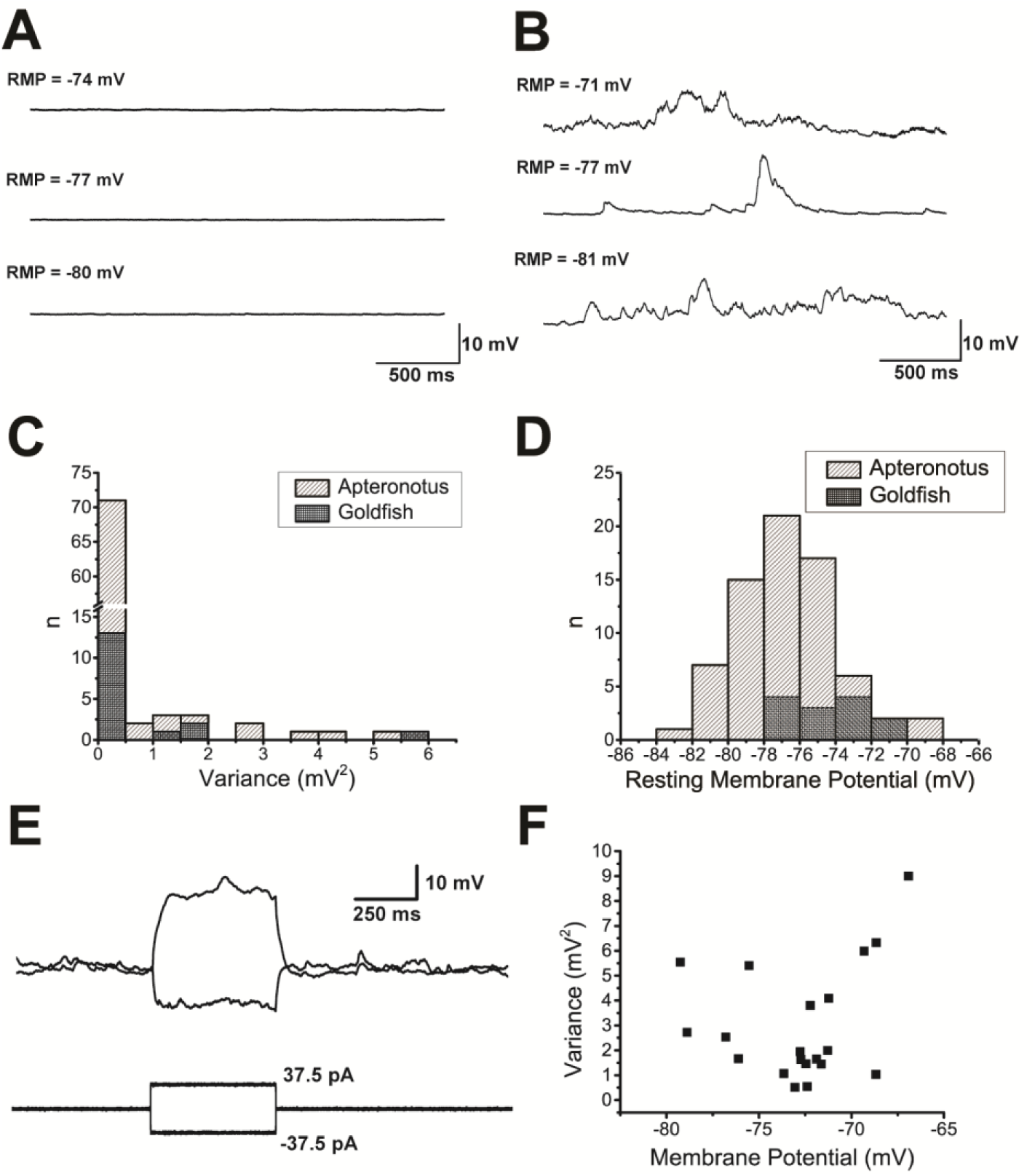
Resting membrane potential of DL neurons. **A**. Three example resting membrane potential (RMP) traces taken from 3 different ‘quiet’ neurons illustrate the membrane potential at which these cells would normally stabilize at naturally (*i.e.* no holding current was applied). The RMP at the start of the recording is shown above each trace. **B**. Three example RMP traces taken from three different ‘noisy’ neurons in which no holding current was applied. In contrast to the ‘quiet cells,’ these cells exhibited strong membrane fluctuations even when they had stabilized at a hyperpolarized potential. **C.** A histogram of the RMP variance for *Apteronotus* (grey) and goldfish (black) DL neurons showing that most neurons were of the ‘quiet’ type where ‘n’ is the number of individual 10 s recording traces that were recorded from all cells. (*Apteronotus*, N = 29 cells; goldfish, N = 7 cells; total n = 85 recordings). **D.** A histogram of the natural RMPs in both the *Apteronotus* and in the goldfish illustrating that the average resting membrane potential of DL neurons is around −77 mV in *Apteronotus* and around −73 mV in goldfish (*Apteronotus*, N = 35 cells; goldfish, N = 11 cells; total n = 71 recordings). **E**. A noisy DL neuron’s response to the injection of ± 37.5 pA current steps in *Apteronotus*, illustrating that the membrane fluctuations are invariant to the membrane potential of the cell. **F.** A scatter plot of the variance and membrane potential, including all recordings (black dots) that had a variance value above 0.5 mV^2^ (*Apteronotus*, N = 6 cells; goldfish; N = 4 cells; total n = 20 recordings).

### Noisy cells

The ‘noisy’ electrophysiological feature has previously been observed in pyramidal cells in the *Apteronotus* hindbrain electrosensory lobe (ELL) and has been attributed to the stochastic opening of voltage-gated ion channels, an effect which becomes stronger as the membrane potential increases toward threshold (Marcoux et al., 2016). We therefore wondered whether noisy DL cells shared these features. DL neurons displayed an *in vitro* RMPs that were relatively more hyperpolarized (*Apteronotus*: −70 mV to −84 mV; goldfish: −66 mV to −78 mV; Fig 2D), compared to the ELL pyramidal cells (−67.8 ± 5.7 mV, Berman and Maler, 1998a) and neither subthreshold depolarizing, nor hyperpolarizing current steps altered the noise fluctuations of *Apteronotus* DL cells (N = 3 noisy cells; Fig. 2E). Additionally, we found that a more depolarized RMP of these noisy cells (*Apteronotus*, N = 6 cells; goldfish, N = 4 cells) was not associated with an increase in noise variance (Fig. 2F). The intrinsic membrane noise of ELL pyramidal cells was shown to unaffected by 10 mM kynurenic acid, an AMPA/NMDA receptor antagonist (Marcoux et al., 2016); this is expected given the lack of recurrent connections in ELL (Maler, 1979; Maler et al., 1981). In contrast, kynurenic acid application (10 mM) completely blocked the membrane potential fluctuations of *Apteronotus* DL cells (N = 2, data not shown).

In some ‘noisy’ cells, spontaneous membrane fluctuations could summate to cause a more sustained depolarization (Fig. 3A). The summating fluctuations were usually between 10 and 20 mV in amplitude and often induced spontaneous action potentials as the membrane potential crossed the spike threshold. The duration of these spontaneous events was estimated to be 425.5 ± 42.4 ms (N = 4 cells), and could reach as long as 800 ms in instances where spontaneous bursting occurred (Fig. 3B). We hypothesize that these events are caused by the summation of multiple postsynaptic potentials, as highlighted by the arrows in Fig. 3C. Based on these observations, we suggest that the membrane noise and spiking are not generated by intrinsic DL cell conductances, but are instead due to synaptic bombardment from neighboring cells within the DL recurrent network (Trinh et al., 2016). In our slice preparation, DL is disconnected from all extrinsic input (Giassi et al., 2012c; Giassi et al., 2012a). As such, the synaptic noise we observed in a subset of DL neurons provides evidence that the activity of the DL recurrent network alone can drive weak spiking activity. We do not currently know why only some neurons show pronounced membrane potential fluctuations.

**Figure 3.**
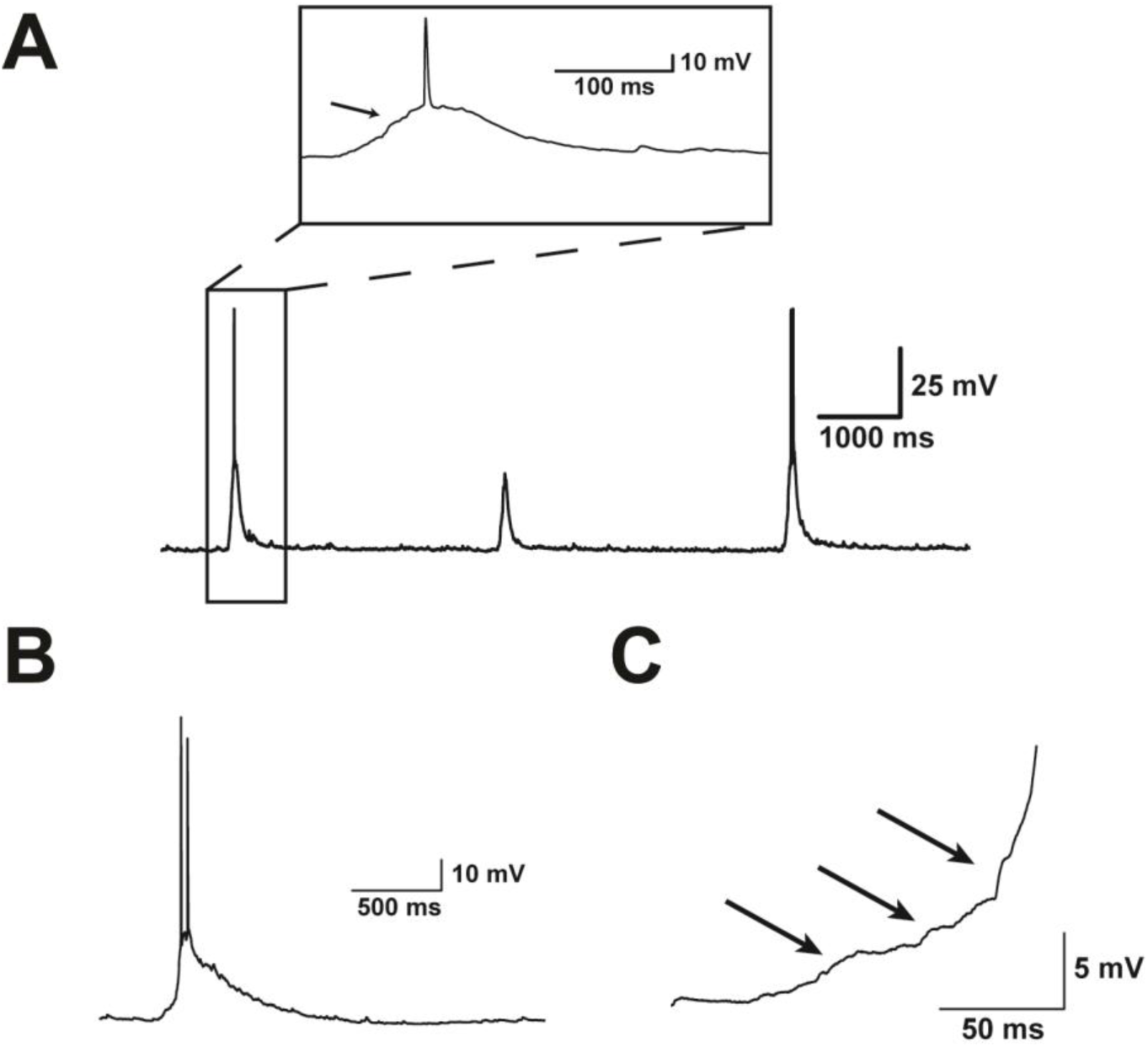
Noisy cells. **A.** Example recording trace from a noisy cell displaying spontaneous membrane potential fluctuations. These fluctuations often vary in size but are usually in the range of several millivolts and can trigger action potentials (spikes), as highlighted by the box showing a magnified version of the first fluctuation. The arrow within the magnified box highlights an example of the small fluctuations that precede spiking. **B.** Example trace illustrating a spontaneous membrane fluctuation that lasted 865.5 ms and produced a short burst of 2 action potentials. **C.** A higher magnification of the rise phase of the spontaneous fluctuation shown in panel 3B. The arrows denote small membrane potential fluctuations that appear to summate, giving rise to a sustained depolarization and spiking.

### Quiet Cells

#### Resting membrane potential, spike threshold and spike discharge patterns

The RMPs of quiet *Apteronotus* DL cells were approximately Gaussian distributed with a mean of −76.7 ± 0.3 mV (N = 29 cells, Fig. 2D), similar to that of goldfish (−74.4 ± 0.7 mV, N = 7 cells). Using the hyperpolarized responses to negative current steps, we calculated an average membrane time constant of 10.28 ± 0.24 ms for these neurons.

We next injected positive current steps in order to generate spiking. An example recording is shown in Fig. 4Ai, illustrating a typical DL neuron response in *Apteronotus*. The same response and spiking pattern was found in all cells regardless of their location within the *Apteronotus* DL region and was also observed in the goldfish DL (Fig. 4B). DL neurons exhibited very pronounced rectification: the membrane potential deflection in response to depolarizing current injections was far stronger than for hyperpolarizing currents of the same magnitude (Fig. 4Aii). This asymmetry is quantified below. In addition, we never observed any ‘sags’ in the response of DL neurons to hyperpolarizing current injections, suggesting that they do not express hyperpolarization-activated cation channels (I_h_).

**Figure 4.**
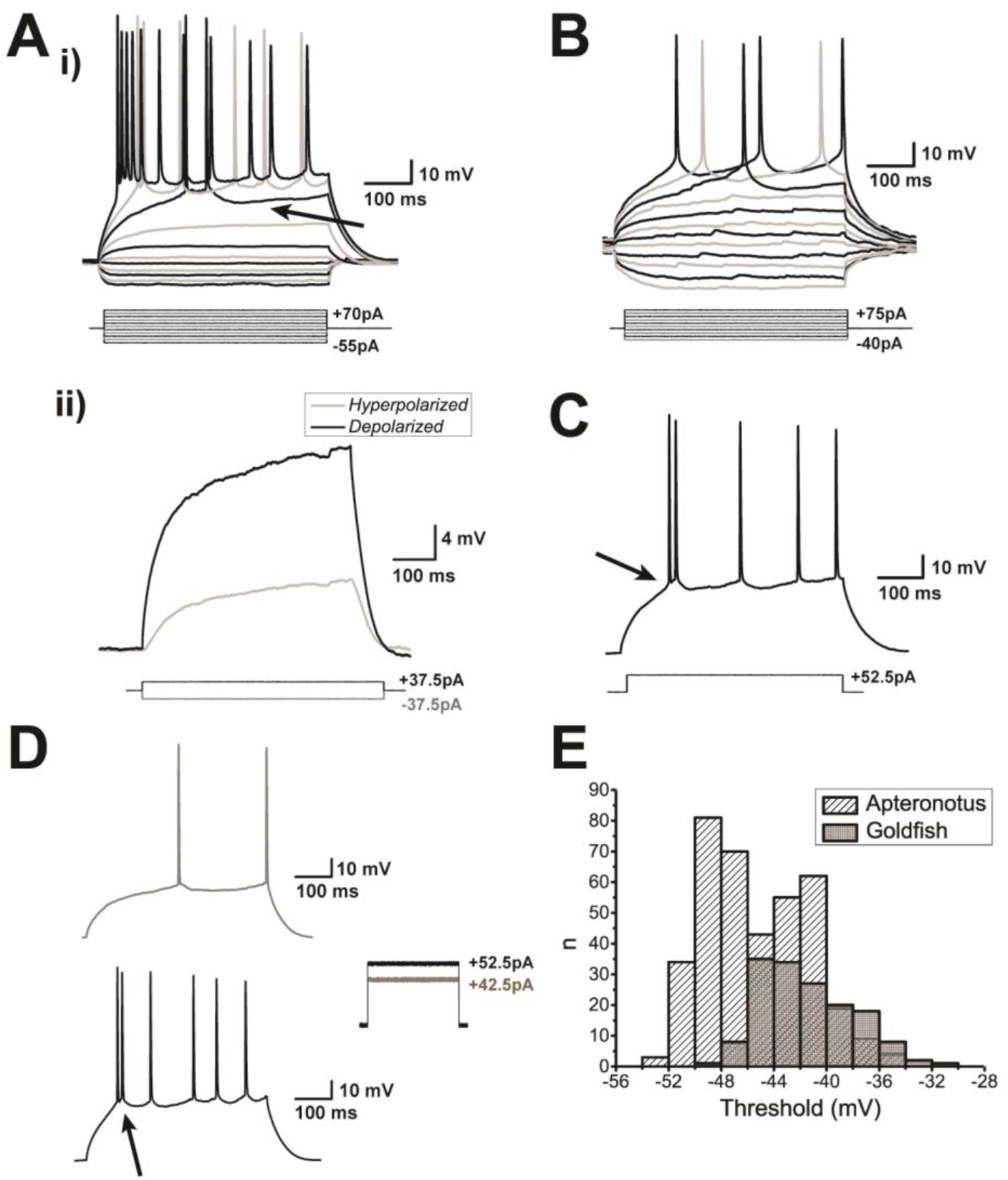
Spiking characteristics of DL neurons. **A. (i)** Example of an *Apteronotus* DL quiet neuron response to the injection of ±500 ms current-steps with varying amplitudes as shown below the response traces. The latency to the first evoked spike clearly decreases with increasing current intensities. However, even at elevated current injections (+70 pA), these cells cannot be driven to a high firing rate (maximum in this case was 22 Hz). This appears to be due, at least in part, to the prominent AHPs that follow the spikes (arrow). There is a large difference between the membrane potential responses to depolarizing versus hyperpolarizing current steps – much stronger responses are seen to positive current pulses. **(ii)** We illustrate this asymmetry by superimposing the absolute responses to equal intensity injections of a hyperpolarizing and subthreshold depolarizing current steps; the response to the hyperpolarizing step is inverted for a clear comparison. DL neuron recordings in goldfish also yielded a similar asymmetry and spiking patterns (data not shown, but see panel 4B). **B**. Example of a goldfish dorsal DL (DLd) neuron response to a standard 500 ms current step injection; the region chosen for these recordings receive inputs from PG similar to the DL neurons in *Apteronotus.*. The responses of these cells were very similar to those of *Apteronotus* DL neurons. **C**. Example recording of a DL neuron in response to a single current step injection. The arrow highlights the location of the threshold for these neurons (see Panel E). **D**. **(i)** A single spike is evoked for currents near spike threshold. **(ii)** After current injections that induce depolarizations exceeding the spike threshold, DL neurons emit a short doublet or triplet burst of spikes at a shorter latency (arrow, *Apteronotus* recording; similar behaviour was seen in goldfish DL neurons). Note that spike amplitude drops slightly but progressively in the 4C and 4D traces. **E**. Histogram of the average threshold of the first current-evoked spike in DL neurons. The spike threshold, which was found using the first derivative of the membrane potential, was ∼ −45 mV in *Apteronotus* and ∼ −42 mV in goldfish. The total number of spikes across all cells used for these estimates was n = 380 in *Apteronotus* and n = 154 in goldfish.

DL neurons discharge very few action potentials (Figs. 4A, B) and the average injected current necessary to reach spike threshold (rheobase) was 38.17 ± 2.52 pA (N =15 cells). Strong current injections (70 pA) only resulted in average firing rates of 15.3 ± 2.4 Hz (N = 15 cells). We defined the spike threshold as the voltage corresponding to a pre-determined fraction of the maximal peak of the first derivative of the membrane potential response to current steps (Azouz and Gray, 2000; methods). Strong current injection in *Apteronotus* DL neurons typically results in an initial high frequency burst of 2 or 3 spikes, followed by an irregular series of spikes separated by AHPs of varying amplitude and duration (Figs. 4C, D); the same pattern was also observed in the DL of goldfish (Fig. 4B and 5A). In *Apteronotus*, the threshold for the first spike is distributed with a mean of −45.3 ± 0.2 mV (N = 22 cells) and has a high degree of overlap with the observed spike threshold for goldfish DL cells (mean: −41.5 ± 0.3 mV, N = 14 cells; Fig. 4E). We measured the mean spike peak amplitude from both the membrane potential at spike threshold (*Apteronotus*: 66.2 ± 1.0 mV, N = 22 cells; goldfish: 50.8 ± 1.0 mV, N = 14 cells) and from the RMP (*Apteronotus*, 95.9 ± 0.5 mV; goldfish, 90.6 ± 0.5 mV). Lastly, we also measured the spike half-width at half-maximum (*Apteronotus*: 2.3 ± 0.1 ms; goldfish: 3.7 ± 0.3 ms).

In summary, the core biophysical properties of DL cells receiving PG input in *Apteronotus* and goldfish (dorsal DL, non-olfactory, Northcutt, 2006; Yamamoto and Ito, 2005) were similar - DL neurons have a hyperpolarized RMP and a high spike threshold and spike only sparsely in response to even strong current injection.

#### Asymmetric input resistance

A striking property of *Apteronotus* and goldfish DL cells is an asymmetry in their response to hyperpolarizing versus depolarizing current steps (Fig.5Ai). In ELL pyramidal cells, an equivalent, though far smaller asymmetry is caused by a persistent Na^+^ channel (Turner et al., 1994) that amplifies excitatory synaptic input (Berman et al., 2001). We tested this possibility by blocking the sodium channels of DL neurons with a local application of 20 µM TTX (control: N = 18 cells, TTX: N = 6 cells). As expected, spike discharge at the previous threshold (∼−45 mV) was completely blocked by TTX (Fig. 5Aii); the small high-threshold spikes evoked with much stronger current injections will be discussed below (Fig. 5Aii, Cii). Upon closer inspection of the neurons’ response to positive current injections, we found that application of TTX did not dramatically change their depolarizing ramp response to peri-threshold current injection (Fig. 5Bi) and, in some cases, would even slightly increase the neuron’s response to positive current injections (Fig. 5Bii). This data indicates that low threshold persistent sodium channels are likely not (or only weakly) expressed in DL neurons.

**Figure 5.**
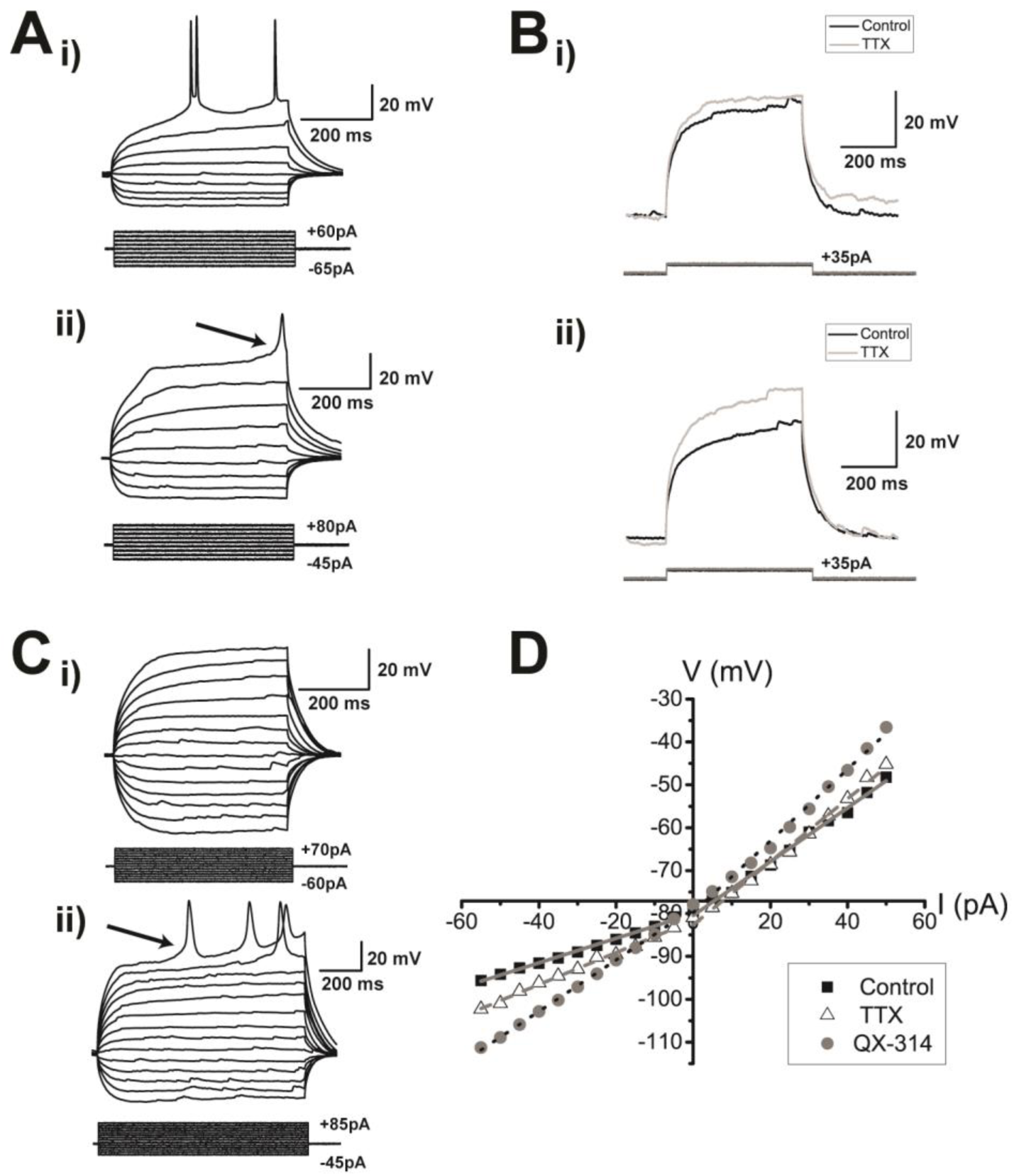
Pharmacological block of sodium and other channels in DL neurons. **A**. **(i)** This panel illustrates a goldfish DL neuron’s membrane potential response to 500 ms current step injections. For +60 pA, large spikes are evoked at a −38.4 mV threshold; in this example, the first spike has a height of 47.5 mV from the threshold and has a half-width of 3.1 ms. **(ii)** The bottom panel shows the responses after bath application of 20 µM TTX, which completely eliminates the large fast spikes. Delayed, broad spikes (amplitude: 22.7 mV from the threshold, half width: 10.5 ms) are now evoked at elevated current levels (+80 pA) with a spike threshold of −21.0 mV. The arrow indicates the approximate location of the threshold for the broad TTX insensitive spike. **B**. **(i)** The response of a DL neuron to a current step at the subthreshold membrane potential before (black) and after (grey) application of TTX. After TTX treatment, the membrane potential did not dramatically change compared to control and in some cases **(ii)** the subthreshold membrane potential was even more depolarized than in the control condition. **C. (i)** Response of an *Apteronotus* DL neuron to current injection steps following QX-314 application via the recording pipette. Fast Na^+^ spikes are eliminated by this treatment, even with strong current injections (+70 pA, 500 ms) that would always evoke spiking in control neurons. **(ii)** Stronger current injection (+85 pA, 1000 ms), evoked delayed broad spikes (amplitude from the threshold = 36.9 mV, half-width; 25.9 ms) with a higher threshold (−8.3 mV) compared to the TTX-insensitive spikes illustrated in panel **A (i)**. Stronger current injections (+85 pA, 1000 ms) evoked several putative Ca^2+^ spikes with a shorter latency to the first spike. The arrow highlights the approximate location of the threshold of the broad Ca^2+^ spike. **D**. Average I-V relationship obtained from subthreshold *Apteronotus* and goldfish DL recordings without the application of any pharmacological blockers (black squares), after the application of 20 µM TTX (white triangles), and with the inclusion of QX-314 within the patch pipette solution (white circles). Both the curves for control and TTX are piecewise linear with the slope being markedly smaller for hyperpolarizing (control; 0.28 ± 0.02 mV/pA, TTX; 0.32 ± 0.04 mV/pA) compared to depolarizing steps (control; 0.69 ± 0.03 mV/pA, TTX; 0.74 ± 0.03 mV/pA). In contrast, the addition of QX-314 has linearized the I-V curve (hyperpolarizing slope = 0.60 ± 0.11 mV/pA, depolarizing slope = 0.86 ± 0.12 mV/pA) with its main effect on the response to hyperpolarizing current injections (see Table 1).

We next plotted the average I-V curves for negative and positive (subthreshold) current injection (*Apteronotus* and goldfish, Fig. 5D). The stronger response to positive versus negative current injection can be clearly seen in the rectification of the I-V curve for the control condition. These curves can be used to compute separate input resistances for positive and negative current injections. Typically, the response to hyperpolarizing current injection is assumed to reflect the passive properties of a neuron and is reported as its input resistance (e.g., ELL pyramidal cells; Mathieson and Maler, 1988; Berman et al., 1997). In DL cells, the input resistance for depolarizing current injection is approximately double that for hyperpolarizing current injection when compared under both control and TTX conditions (Table 1; paired t-test; control; p = 3.3×10^−12^, TTX; p = 9.9×10^−6^). The addition of TTX had no significant effect on the hyperpolarizing slope (one way ANOVA; p = 0.32), nor did it have any significant effect on the input resistance for the depolarizing slope (Table 1: one way ANOVA; p = 0.42; Fig. 5D). Thus, it appears that there is no contribution of persistent Na^+^ channels to the resting membrane potential of DL neurons, in accordance with the small effects of TTX observed in Fig. 5B.

**Table 1:** I-V slope measurements obtained from the depolarizing and hyperpolarizing responses of DL neurons in both teleost species for both the TTX and QX-314 experiments.

| Conditions | Depolarizing Slope<br>(GΩ) | Hyperpolarizing Slope<br>(GΩ) |
| --- | --- | --- |
| Control<br>(N = 18 cells) | 0.69 ± 0.03 | 0.28 ± 0.02 |
| TTX<br>(N = 6 cells) | 0.74 ± 0.03 | 0.32 ± 0.04 |
| QX-314<br>(N = 6 cells) | 0.86 ± 0.12 | 0.60 ± 0.11 |

To further investigate the basis of the observed asymmetrical response to current injection, we have also recorded DL neurons using an intracellular solution containing 5 mM QX-314, a blocker of Na^+^ channels, as well as some K^+^ and Ca^2+^ channels (Talbot and Sayer, 1996) (Fig. 5C, D; control, N = 18 cells; QX-314, N= 6 cells). QX-314 has previously been used to block all Na^+^ channels in *Apteronotus* ELL pyramidal cells (Berman et al., 2001). The I-V graph constructed from the QX-314 experiments showed a higher depolarizing versus hyperpolarizing input resistance (paired t-test; p = 2.3×10^−4^), similar to control and TTX conditions (above). There was a small increase in input resistance for the depolarizing current injection that failed to reach significance (Table 1: one way ANOVA, p = 0.07; Fig. 5D). In contrast, there was a large and highly significant increase of input resistance in the responses to hyperpolarizing current injections – it more than doubled over control values (Table 1: one way ANOVA, p = 5.9×10^−5^). Since we only expect K^+^ permeating channels to be open at such hyperpolarized membrane potential, we attribute this effect to the ‘non-specific’ actions of QX-314 (Perkins and Wong, 1995; Slesinger, 2001). The results of the TTX and QX-314 experiments lead to two hypotheses: first, the subthreshold response of DL cells to depolarizing input is mainly due to their passive membrane properties. Second, the RMP of hyperpolarized DL cells is likely due to a strong rectifying K^+^ conductance that is blocked by QX-314 and typically prevents the cell from deviating from the reversal potential of K^+^ ions (Fig. 5). Given that GIRK channels are ubiquitous in the mammalian cortex (Luscher and Slesinger, 2010; Lujan and Aguado, 2015) and can be blocked by QX-314 (Zhou et al., 2001), we suspected them to also be present in the teleost pallium. To confirm the presence of GIRKs in DL, we used a RT-PCR approach to show the expression of GIRK channels in different brain regions (DL, subpallium, tectum/torus, cerebellum, ELL and hindbrain) using a primer pair hybridizing in conserved segments of all GIRK paralogs. Unsurprisingly, pan-GIRK amplicons were found in all brain regions, but were not present in the control (Fig. S1) suggesting that GIRK channels are ubiquitously expressed in the *Apteronotus* brain.

#### Voltage dependent calcium conductance

In the presence of TTX, strong current injections (>80 pA) were able to evoke a broad (half-width: 33.0 ± 3.1ms) spike with a very high threshold (mean threshold: −21.2 ± 0.5mV, N = 4 of 6 cells; Fig. 5Aii). Spike amplitude was 18.6 ± 0.7 mV from the threshold potential and 79.1 ± 0.9 mV from the RMP. Similar to the TTX results, QX-314 treated cells did not produce any action potentials at the threshold for control cells (Fig. 5Ci), but did produce broad spikes at much higher stimulus intensities (spike half-width; 21.1 ± 1.1 ms; height = 31.3 ± 0.7 mV from threshold and 100.6 ± 0.9 mV from RMP, N = 4 of 6 cells, Fig. 5Cii). The average threshold for these broad spikes was found to be at −6.8 ± 1.3 mV, which is also consistent with the range of voltages that has been reported for the activation of HVA Ca^2+^ channels (Tsien et al., 1988). Therefore, we hypothesize that DL neurons express HVA Ca^2+^ channels that will likely be activated by Na^+^ mediated action potentials.

#### After-hyperpolarizing potentials

DL neurons exhibit a strong AHP (Figs. 4-6). We have previously shown that DL cells express both SK1 and SK2 channels (Ellis et al., 2008) and that UCL1684 is highly effective at blocking such channels (Harvey-Girard and Maler, 2013). We therefore bath-applied 30 µM UCL1684, resulting in a significantly diminished AHP compared to the control conditions (Fig. 6A). To quantify this AHP reduction, we measured the AHP amplitude (Fig. 6Ci) and the area under the AHP (Fig. 6Cii) following the first single spike obtained in response to current injection. The addition of UCL1684 reduced the average amplitude of the first AHP to half its control value (Fig. 6Ci: control: 3.5 ± 0.3 mV, N = 13 cells; UCL1684: 1.4 ± 0.2 mV; N = 7 cells; two sample t-test; p = 0.0003). A similar reduction was also observed when comparing the area under AHPs: from 1980.3 ± 192.6 mV·ms to 701.4 ± 128.4 mV·ms (Fig. 6Cii, two-sample t-test; p = 0.0002). In contrast, after the addition of the SK channel agonist EBIO (1 mM; Ellis et al., 2007), current injection evoked very few spikes; thus current steps were increased to 1000 ms. As expected, the average AHP amplitude increased from 3.5 ± 0.3 mV to 6.7 ± 1.0 mV (control, N = 13 cells; EBIO, N = 6 cells; two sample t-test; p = 0.001), while the area under the curve also increased from 1980.3 ± 192.6 mV·ms to 3952.2 ± 277.5 mV·ms (two sample t-test, p = 0.00002).

**Figure 6.**
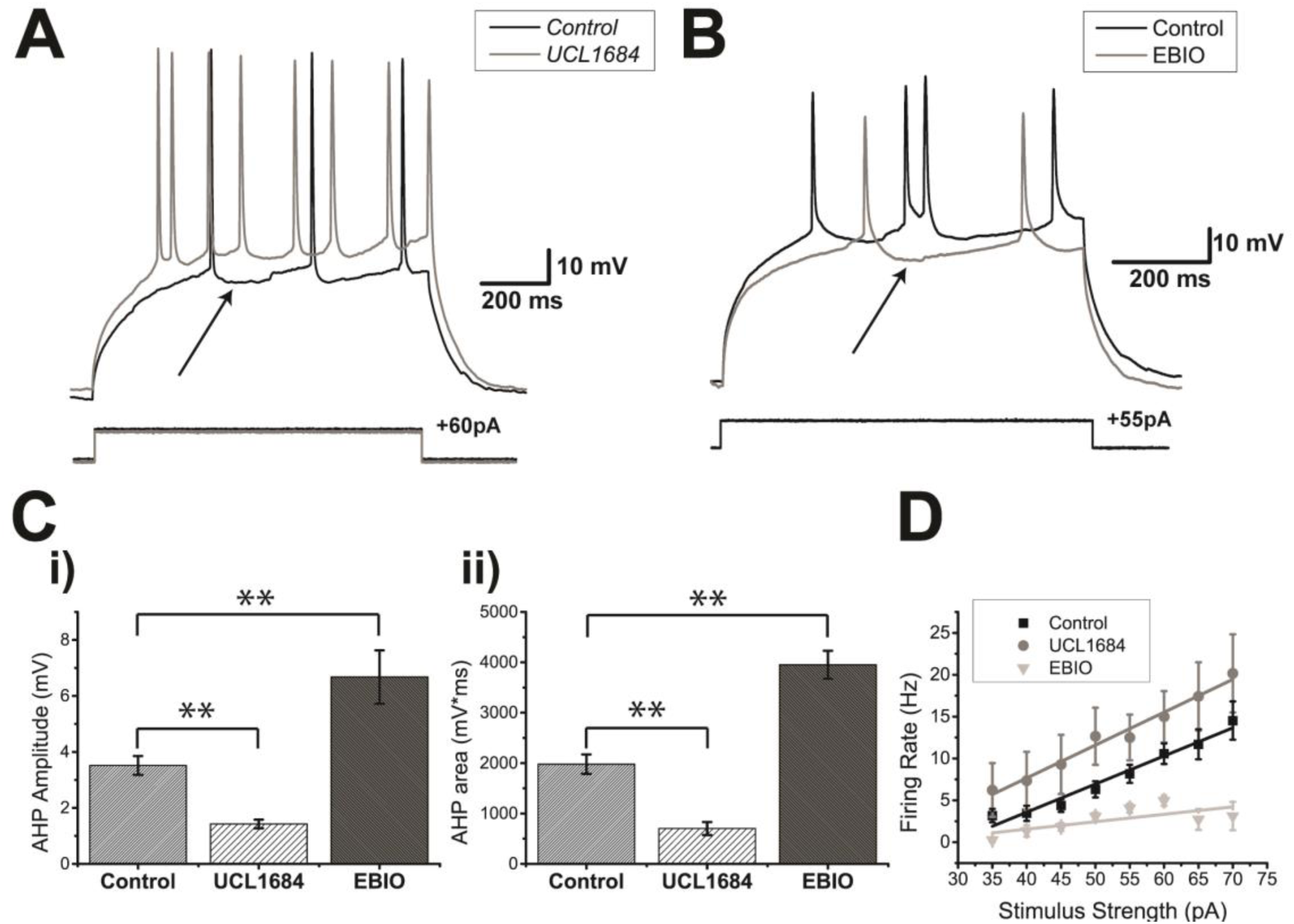
SK-mediated potassium channels contribute to the AHP of DL neurons. **A**. *Apteronotus* DL neuron response to 500 ms step current injection before (black trace) and after the bath application of 30 µM UCL1684 (grey trace). The black arrow shows the minimum membrane potential between two spikes and is used to estimate the amplitude of the AHP by comparison with the membrane potential immediately preceding the first action potential. The prominent AHPs seen in the control condition are reduced by this treatment and the spike rate has also increased (from 3 spikes to 8 spikes). **B**. DL neuron response to the injection of 1000 ms current steps before (black trace) and after bath application of 1 mM EBIO in the goldfish (grey trace; a longer pulse was needed in order to increase the likelihood of evoking more than one spike). The amplitude of the AHP (arrow) was increased by this treatment and the spike rate has been reduced (from 4 to 2 Hz). **C.** Average amplitude **(i)** and average area under the membrane potential **(ii)** of the AHP following the first spike of DL neurons in response to current steps (control, N = 12 cells; UCL1684, N = 7 cells; EBIO, N = 5 cells). Both the amplitude and the area under the AHP are significantly diminished after the application of UCL1684, while a strong increase was observed after the application of EBIO. **D**. Average firing rate plotted as a function of the amount of current injected for the control condition (black trace), the UCL1684 condition (grey trace) and the EBIO condition (light grey trace) in both *Apteronotus* and goldfish (control, N = 28 cells; UCL1684, N = 7 cells; EBIO, N = 5 cells). The firing rate increases for all current injections after UCL1684 application, while the firing rate decreases after the EBIO application.

Blocking SK channels also increased the current-evoked firing rate compared to the control condition (Fig. 6D; control, N = 28 cells; UCL1684, N = 7 cells; two-way ANOVA; p = 0.0013), while EBIO reduced the evoked firing rate since the cell required a longer time to reach spike threshold after the first spike (Fig. 6B, D; N = 6 cells; two-way ANOVA; p = 0.000092). We conclude that the SK1/2 channels of DL neurons act to promote spike frequency adaptation that acts as negative feedback on their firing rate in response to excitatory input.

Finally, we wanted to confirm whether SK channel activation in DL neurons could be blocked by preventing Ca^2+^ activation of the channel. We recorded DL neurons in *Apteronotus* using an intracellular solution containing 10 mM BAPTA, a Ca^2+^ chelator (N = 7 cells; Fig. 7A). In all cases, the AHP was completely abolished, unlike the partial AHP block obtained with UCL1684. This suggests that another unidentified Ca^2+^ activated K^+^ channel may also be contributing to the AHP. Further work will be required to investigate this possibility. The firing rate also dramatically increased compared to the control condition (control, N = 28; UCL1684, N = 7; BAPTA, N = 7 cells; Fig. 7B; two-way ANOVA; p = 1.5 × 10^−15^) and compared to UCL1684 treatment (two-way ANOVA; p = 0.00063). Furthermore, this BAPTA-induced increase in firing rate was also accompanied by a significant reduction in spike height compared to both control (Fig. 7C; two-way ANOVA; p = 2.1 × 10^−12^) and UCL1684 conditions (two-way ANOVA; p = 1.6 × 10^−6^). In contrast, the difference in spike height between the UCL1684 and control did not yield a significant difference (two-way ANOVA; p = 0.14). We hypothesize that Na^+^ channel inactivation may be causing this reduction (see below). Another distinctive feature of the DL neuron’s spiking response during the BAPTA application was the increase in spike width occurring along successive spikes and typically becoming most prominent by the 8^th^ spike (Fig. 7A, D). In the control and UCL1684 conditions, there was a slight increase in spike width that might be linked to the inactivation of SK channels. However, in the BAPTA condition, the spike width increased dramatically with successive spikes (Fig. 7A, 7D) compared to control (two-way ANOVA; p = 1.3 × 10^−31^) and UCL1684 conditions (two-way ANOVA; p = 3.7 × 10^−10^). In contrast, the difference between the control and BAPTA conditions was not significant up until the third spike (two-way ANOVA; p = 0.24), suggesting that the spike width increase is caused by a cumulative process. Calcium channels typically inactivate via a Ca^2+^-dependent mechanism (Simms and Zamponi, 2014), leading us to hypothesize that this dramatic change in spike width may be caused by a decrease in Ca^+^-dependent inactivation of the Ca^2+^ channel leading to an increase of its open time (see Discussion).

**Figure 7.**
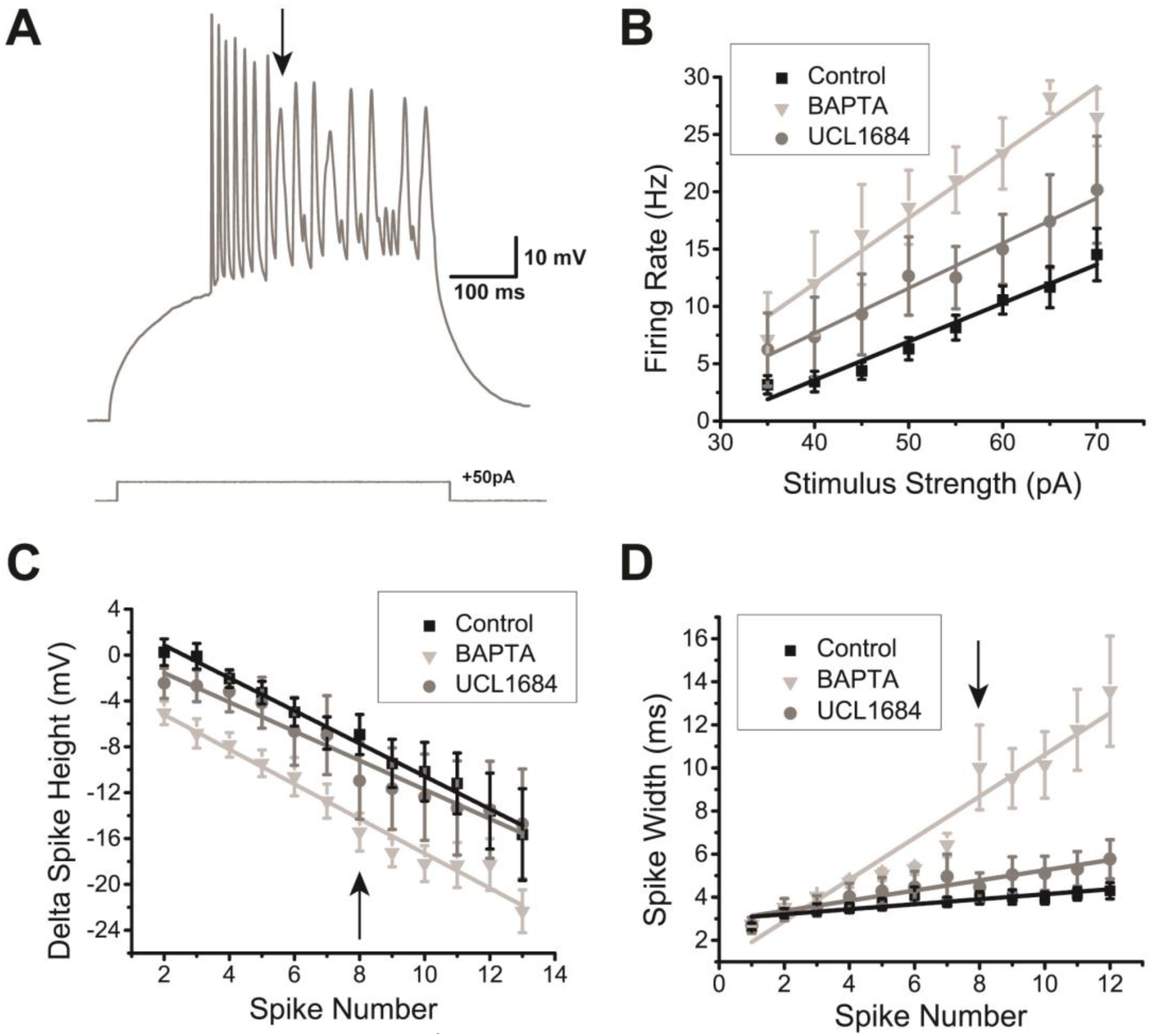
The effect of intracellular Ca^2+^ chelation on DL neuron responses to depolarization. **A**. Example recording trace (*Apteronotus*) with 10 µM BAPTA added to the internal solution of the patch pipette. The AHP appears to be completely eliminated which promotes higher frequency spiking; note that successive spike heights drop continuously for the first seven spikes. By the 8^th^ spike, very prominent spike broadening begins and the spike height drops to an even greater degree compared to the UCL1684 application in Fig. 6B. **B.** The average firing rate was plotted as a function of the amount of current injected for the control (black trace, N = 28 cells), UCL1684 (grey trace, N = 7 cells) and BAPTA conditions (light grey trace, N = 7 cells). The addition of intracellular BAPTA promotes an even stronger increase in firing rate compared to the addition of the SK channel blocker UCL1684. **C.** The average difference in spike height between the n^th^ spike and the first spike was plotted as function of successive spikes obtained after a 500 ms current step injection for all 3 conditions mentioned in 7B. The addition of UCL1684 did not strongly affect the spike height, unlike the addition of BAPTA, which reduced the spike height across successive spikes following a step current injection. The arrow highlights the 8^th^ spike, which marks the beginning of the non-linearity in the BAPTA condition. **D.** The average spike width was plotted as a function of successive spikes, similar to panel 7C.UCL1684 application has only a minimal effect on spike width. The presence of intracellular BAPTA increased the spike width across successive spikes during a step current injection when compared to the other conditions. The arrow indicating the 8^th^ spike marks a strong change in spike width, as denoted by the arrow in 7A.

#### Dynamic AHP and spike threshold

Although the presence of the AHP greatly reduces the firing rate, we also observed that after successive spikes, the AHP itself decreased (Fig. S2A) and the spike threshold increased (Fig. 8A). To better quantify the AHP modulation, we measured the difference in AHP amplitude between the first two spikes of a current-evoked spike train that did not show an initial burst. We found that there was a significant reduction in AHP amplitude that was invariant to the time length of the AHP (Fig. S2B black squares; N= 26 cells; one-sample t-test, p = 4.45×10^−27^). For recording traces that showed initial bursts, we examined the first spike pair following the burst and found a similar reduction in AHP (Fig. S2B, grey triangles; N = 20 cells; one sample t-test, p = 1.26×10^−13^). This reduction is presumably caused by Ca^2+^-induced inactivation of the HVA Ca^2+^ channels, which will decrease the total amount of Ca^2+^ available to the cell and limit the activation of SK channels.

**Figure 8.**
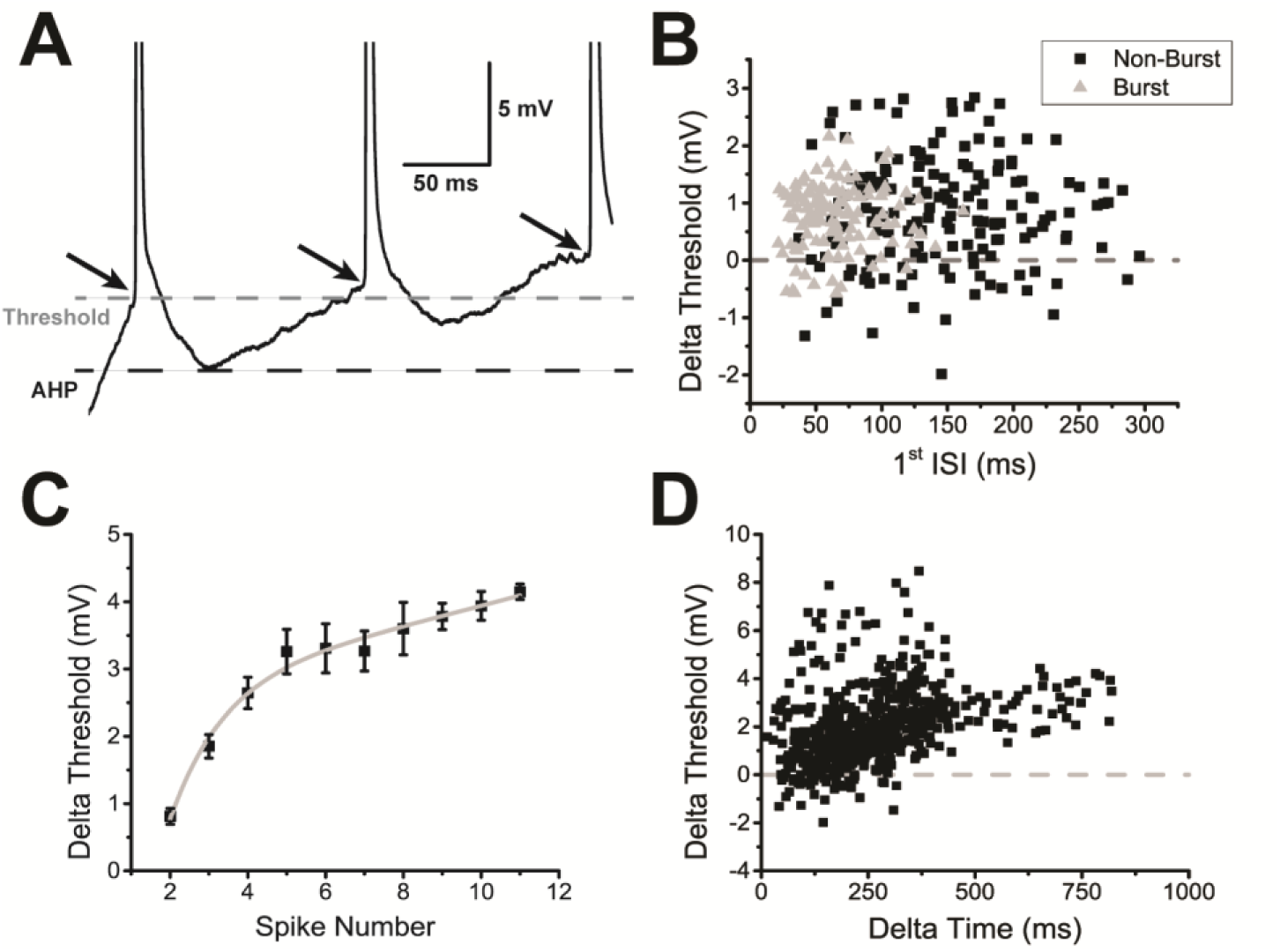
DL neuron spiking causes a decrease in AHP amplitude and an increase in spike threshold. **A.** A magnified view of the first 3 spikes in an example trace of a DL neuron’s response to a +60pA current injection where the black dashed line is placed to coincide with the minimum of the first spike’s AHP and the grey dashed line is placed to coincide with the first spike’s threshold. The black arrows highlight the progressive increase in spike threshold following consecutives spikes. **B**. The increase in spike threshold between the second and first spikes was plotted in the same manner as a function of the first interspike interval (ISI). Individual black squares represent a pair of spikes that were taken from a trace which did not contain an initial burst (total of 160 non-burst pairs), while individual triangles represent a pair of spikes that were taken from a trace displaying an initial burst of spikes as in Fig. 4C (total of 117 burst spike pairs). The majority of the spike thresholds increased (over 300 ms) with no evident recovery. **C**. The difference in average spike threshold between the n^th^ spike and the first spike is plotted as a function of the spike number. The subsequent curve was fit with a double exponential equation (y = 2.72\**e*^0^.^04x^ −9.0*e*^−0^.^72x^; R^2^ = 0.987). **D.** The increase in spike threshold between the n^th^ spike and the first spike is plotted as a function of the time interval between them. Each black square represents a spike pair (total of 573 spike pairs). Overall, the increase in threshold appears to be larger following longer timer intervals.

Even with the spiking-induced reduction of the AHP, DL neurons could not surpass a sustained firing rate of 30 Hz (Fig. 7B), which suggests the presence of an additional mechanism(s) that limits firing rate. In ELL pyramidal neurons, spike threshold fatigue has been shown to limit the firing rate whenever a burst occurs (Chacron et al., 2007). Upon closer inspection, we found a significant increase in spike threshold during long spike trains (Fig. 8A). This dynamic spike threshold was also found to be invariant to the inter-spike interval (up to ∼300 ms) for both non-burst traces (Fig. 8B, black squares; one-sample t-test, p = 1.24×10^−22^) and for traces containing an initial burst (Fig. 8B; one sample t-test, p = 8.30×10^−28^). Next, we wanted to confirm whether the threshold fatigue that was observed in DL neurons may be caused by the history of past spikes, *i.e.*, whether Na^+^ channel inactivation due to continuous spiking may influence the spike threshold. We examined the difference in threshold for all non-burst traces to see whether it varies throughout a spike train. The threshold increase continued to at least 10 spikes and could be fitted by a double exponential function (equation: y = 2.72*e*^0^.^04x^ + − 9.0*e*^−0^.^72x^; R^2^ = 0.987; Fig. 8C); here we considered only the number of spikes and not the duration of the spike train. A similar analysis where the difference in threshold was compared to the time between spikes instead of the spike number also led to the same conclusion: this effect became more prominent after long periods of depolarization despite the variability in the number of intervening spikes (Fig. 8D). These results suggest that the increase in threshold is caused by an accumulation of slow Na^+^ channel inactivation.

To quantify the duration of this spike threshold adaptation, we developed a protocol in which a long ramp current (evoking multiple spikes) was injected followed by a shorter ramp current (evoking one spike) at various inter-stimulus time intervals (Fig. 9A, upper panel). This protocol induced spike threshold fatigue during the first current injection, while the second current injection was used to test for time-dependent changes in spike threshold. We found that the increase in spike threshold between the first and second current injection was significantly higher at short compared to longer time intervals (Fig. 9A, bottom panel). Using these changes in threshold, we found that the recovery from this spike threshold fatigue had a highly variable time constant ranging from 300 ms to 900 ms with an average time constant τ_exp_ = 637.28 ± 85.9 ms (Fig. 9B). This suggest that the decrease in cell excitability caused by the dynamic threshold can operate on the timescale of hundreds of milliseconds.

**Figure 9.**
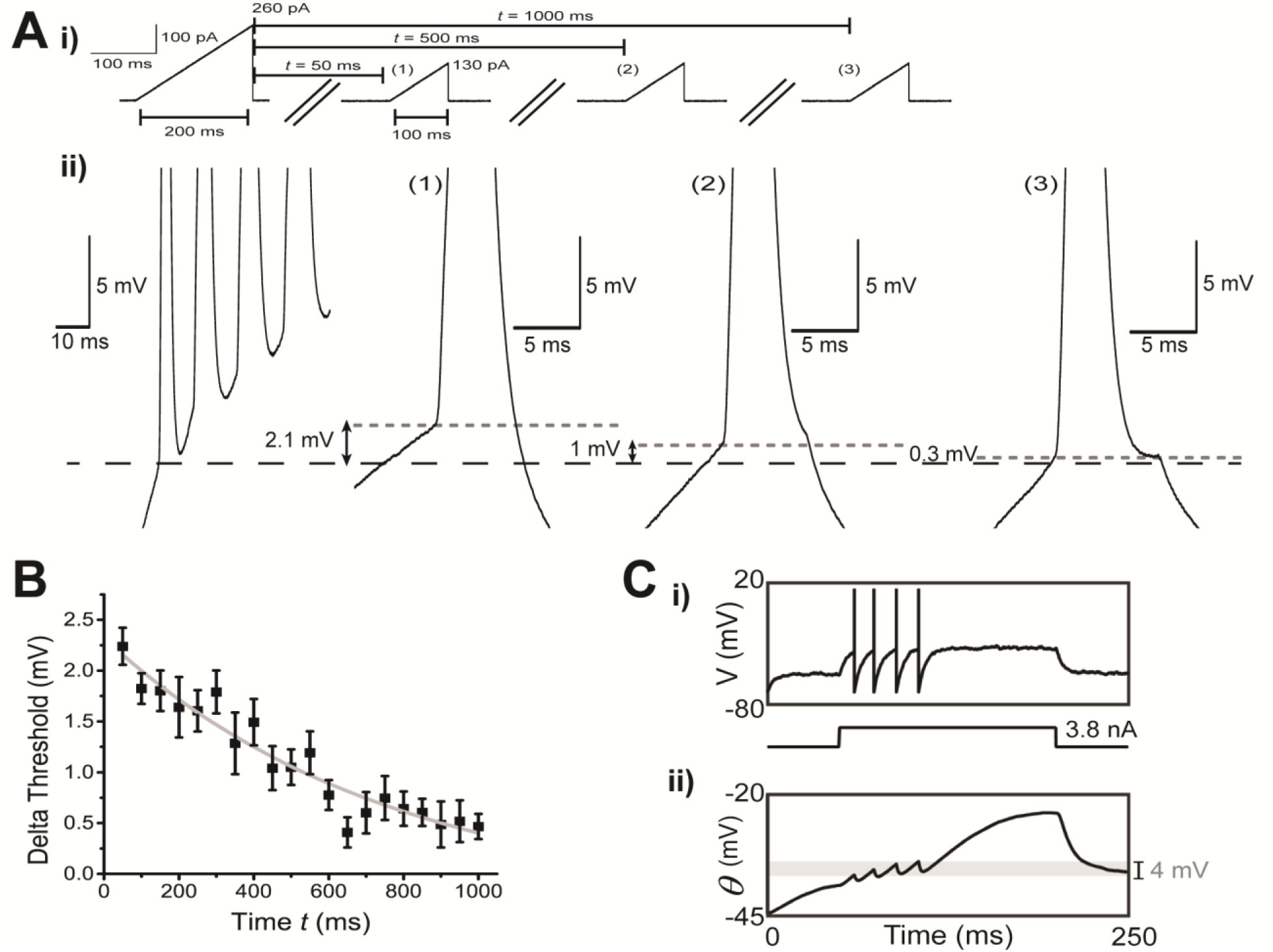
DL neuron spike threshold adaptation can last up to hundreds of milliseconds. **A. (i)** Two ramp current injections separated at various times *t* (50 ms, 500 ms and 1000 ms) were used to measure the time constant of the spike threshold adaptation. Although both ramp current injections have the same slope (slope = 1.3 pA/ms), the objective of the first ramp current was to induce an accumulation of Na^+^ channel inactivation through the firing of multiple action potentials and therefore was stronger than the second ramp current which would only produce one action potential. **(ii)** A magnified view of the example responses obtained after the first ramp current injection and after the second ramp current injections at times *t* = 50 ms, 500 ms and 1000 ms respectively. The black dash line is aligned to the first spike’s threshold obtained after the first ramp current injection while the grey dotted lines are aligned to the spike threshold obtained from the second ramp current injection for the various times *t* mentioned previously. **B.** The average difference in spike threshold (for the first spike only) between the first and second ramp current injections were plotted as a function of the time *t* between each ramp injection. The resulting curve was fitted with an exponential equation (y = 2.38\**e*^−0^.^0017x^, R^2^ = 0.917). **C**. We used a simplified exponential integrate and fire model with fast (_*τ* *f*_ = 15 ms) and slow (*τ* _*s*_ = 500 ms) Na^+^ channel inactivation in an attempt to connect the apparent Na^+^ channel inactivation (Fig. 7A) with the increase in spike threshold over multiple spikes (C, D). **(i)** When driven by a step current, the model produces a small number of spikes at frequencies consistent with the data; however, the neuron quickly ceases discharge despite continuous application of the strong positive current. **(ii)** This result can be easily understood in terms of the dynamic spike threshold (*θ*), which increases because of cumulative slow inactivation of the Na^+^ channel (*h*_*f*_ and *h*_*s*_ not shown). Note that the threshold changes by approximately 4 mV (grey shading) over the course of a few spikes, in line with the upper bound for threshold increases seen between DL cell spikes (panel **i**).

To further investigate whether Na^+^ channel inactivation is responsible for the observed increase in threshold, we employed a model with minimal assumptions: the inactivating exponential integrate and fire model (Platkiewicz and Brette, 2011). This model includes a slow inactivation term, as well as the traditional fast inactivation term associated with Na^+^ channels; inactivation kinetics for both forms were derived from our data (see Methods). Since we were primarily interested in the effect of sodium inactivation on the spike threshold, the minimal model omits AHP dynamics and Ca^2+^ currents (see Methods). We found that the addition of slowly inactivating Na^+^ channels, as suggested by the effects of BAPTA (Fig. 7A), was itself sufficient to qualitatively reproduce the response of DL neurons to current injection and predict an increase in spike threshold that was similar to that observed in our whole-cell recordings (Fig. 9C). We therefore conclude that the accumulation of slow Na^+^ channel inactivation, caused by spike discharge and simple depolarization, may act as a source of negative feedback to reduce the cell’s firing rate via an increase in spike threshold.

#### Modeling DL neurons with a generalized leaky integrate and fire (GIF) model

One aim of our descriptive analysis of DL cell physiology was to develop a simple model of these cells that would permit subsequent modeling of spatiotemporal spiking activity patterns within the putative DL attractor network (Trinh et al., 2016). Recently, a new algorithm has been developed by Pozzorini et al. (2015) for modeling cortical pyramidal cells based on physiological recordings of their response to modulated noise inputs. This method involves the extraction of model parameters and constructing a generalized integrate and fire model (GIF), which best reproduces observed spiking activity. Since the entire noise stimulus protocol lasted for several minutes, we wanted to verify the health of the patched cell by injecting standard 500 ms step currents before and after the protocol, denoted by “pre” and “post” (Fig. 10A); we used the responses to these current steps to ascertain whether the stimulus protocol had changed the spike height and width, as well as to determine if there were any long-term changes in cell excitability (N = 9 of 13 cells, see below). The patched cell undergoes two distinct stimulus phases: the training phase and the test phase (Fig. 10A). During the training phase, the cell is injected with a peri-threshold noise stimulus that should induce spiking at a rate of ∼10 Hz. The resultant response is then used to obtain the parameters necessary to construct the GIF model. In the test phases, the cell is subjected to another peri-threshold noise stimulus but of a shorter duration. One problem with applying this protocol to DL neurons was that we could not safely drive them to a sustained 10 Hz firing rate and still maintain cell viability and we may therefore have under sampled the spiking dynamics (Fig. 10).

Based on spike height measurements taken throughout the pre/post step stimulation, the recorded cells all appeared to be healthy throughout the protocol. Surprisingly, the cell’s response to the peri-threshold stimulus yielded extremely variable firing patterns in both *Apteronotus* (N = 10) and goldfish (N = 3) DL cells; we observed membrane potentials that decreased (4 of 13, Fig. 10B), remained stable (4 of 13 cells, Fig. 10D) or increased (5 of 13, Fig. 10D) through the training recordings during the training phase. The RMP generally remained stable in the rest periods between test stimulation; one exceptional increase is illustrated in Fig. 10B. Despite the general membrane potential stability, the firing rates during test phase stimulation varied erratically, albeit over a small range. Similar to the variation during the training phase, an increase in firing rate was often accompanied by an increase in membrane potential. This variability is not a necessary consequence of the lengthy stimulation protocol. For example, ELL pyramidal cells will maintain consistent discharge rates for over 100 seconds of noise stimulation (Mehaffey et al., 2008). We note that the increases in firing rate during training and test stimulation is contrary to the increases in spike threshold we observed with static stimulus pulses; we had expected to find overall decreases in firing rate in response to the relatively long stimuli and cannot account for this discrepancy.

In some cases, the neuron’s excitability seemed to have changed in the post-protocol period when compared to the pre-protocol period. The average firing rate during the pre-protocol stimulation was 8.2 ± 1.9 Hz, while the average firing rate during the post-protocol stimulation was 12.0 ± 2.4 Hz (N = 9 cells, two sample t-test; p = 0.078). In half of the cases, we observed an increase in excitability that was accompanied by an increased firing rate and a decrease in rheobase current (5 of 9 cells, Fig. 10B,C, Fig. S3A; paired t-test; p = 0.021), while in other cases, the cell’s excitability remained unchanged (4 of 9 cells, Fig. 10D, Fig. S3B; paired t test; p = 0.26). To confirm that this change in excitability was caused by the current injection used in the protocol, we also recorded some control cells where the standard 500 ms step current injection was applied before and after a rest period that was equal in duration to the fitting protocol, *i.e.*, 7 min (N = 8 cells). Unsurprisingly, the firing rate of the post-protocol injection remained unchanged in almost all cases (7 of 8 cells) and the average change in rheobase was also statistically insignificant (Fig. S3C; paired t test; p = 0.44), suggesting that the change in excitability observed after the current injection protocol was not caused by an intrinsic stochastic mechanism. The difference between experimental and control conditions did not, however, reach statistical significance (Chi-square test: 3.438; p = 0.063). This excitability variation may be explained by two, not mutually exclusive mechanisms. For instance, long-lasting discharge (tens of seconds) at even low firing rates (Fig. 10C) might modulate DL cell voltage-or Ca^2+^-gated ion channels, thereby influencing DL cell excitability - modulation might be induced by Ca^2+^-activated signalling cascades. Alternatively, discharge of even a single DL cell might induce synaptic plasticity of its recurrent synapses and affect local network activity. In either case, the nonstationary behaviour of DL cells on the order of tens of seconds makes it difficult to construct simple models that capture their essential dynamics.

To confirm the accuracy of the GIF model, we analyzed the M*_d_ value, which corresponds to a goodness of fit measure computed from the difference between the predicted and experimental spiking activity (Pozzorini et al., 2015). Among the tested cells, only a few had generated acceptable fits. In almost half of the cases, although stimulus induced fluctuations were captured, the GIF model predicted less spikes than were obtained in the experimental recordings (6 out of 13 cells) generating an average M*_d_ value of 0.51 ± 0.05 for all 13 cells in both teleost species. Furthermore, there was no correlation between the goodness of fit of the model and the change in excitability that was observed during the post-protocol stimulus - we observed both good fits and bad fits regardless of the change in excitability. An example good fit for a cell demonstrating an increase in excitability is shown in Fig. 10E (M*_d_ = 0.84), which also corresponds to the example cell shown in Fig. 10B, while an example good fit for a cell which did not change in excitability is illustrated in Fig. 10F (M*_d_ = 0.61). Overall, the spiking pattern of the DL neuron wasn’t sufficiently stable and strong (>10 Hz) during both the training phase and the test phases of the fitting protocol (Fig. 10F). As such, the GIF model was not able to predict the spiking pattern of the majority of the tested cells with high accuracy. Despite their simple morphology (Giassi et al., 2012b), DL neuron dynamics are complex and heterogeneous (Fig. 9B,C; Fig. 10). It will require more detailed physiological studies and more sophisticated models to identify the long timescale properties of DL cell discharge and capture the cellular bases of their heterogeneity.

## Discussion

The work presented here is, to our knowledge, the first study of the biophysical properties of teleost DL neurons. Our previous work mapped local DL circuitry (Trinh et al., 2016), the organization of thalamic and other input to DL (Giassi et al., 2012c), and the telencephalic connectivity of DL (Giassi et al., 2012a; Elliott et al., 2017). We have an accurate analysis of how the electrosensory system contributes to spatial learning (Jun et al., 2016), and have characterized the PG (thalamic) input to DL in response to object motion (Wallach et al., 2018). Although we have yet to examine the intrinsic and extrinsic properties of DL synaptic input, we believe that the constraints imposed by DL circuitry, behavioural function plus recent theoretical analyses, are sufficient to generate testable hypotheses of the computations performed by DL during spatial learning. Below, we first summarize the main conclusions of our work and then discuss whether the biophysical properties of DL neurons and their connectivity are compatible with the critical role of DL in spatial learning and memory. In particular, we suggest that DL neurons possess the minimal requirements to be labelled as sparse coders. Next, we suggest that spike threshold adaptation is key to the extraction of spatial information in DL from the time stamped electrosensory input conveyed by PG (Wallach et al., 2018). Our hypothesis relies on a previous theoretical model of time coding cells (Itskov et al., 2008) that utilizes, as an essential ingredient, spike threshold adaptation with a long recovery time constant.

Our main results show that DL neurons express a combination of ion channels that have been reported for many other types of neurons. DL neurons have a hyperpolarized RMP. We hypothesize that is due, at least in part, to GIRK channels. GIRKs can hyperpolarize neurons by at least 8 mV under basal conditions (Luscher and Slesinger, 2010) and have been shown to set the RMP of dorsal cochlear nucleus neurons to a hyperpolarized level (Ceballos et al., 2016). DL neurons also have a high spike threshold and theoretical analyses suggest this may attributed to a low density of voltage-gated Na^+^ channels (Platkiewicz and Brette, 2010). Furthermore, our results also imply the presence of HVA Ca^2+^ channels, which activate a strong SK channel mediated AHP that strongly reduces current-evoked spiking. We propose that the combination of a hyperpolarized RMP, the low input resistance at hyperpolarized potentials (Table 1), a high spike threshold and strong AHPs will greatly reduce DL cell excitability and therefore prevent incoming excitatory synaptic input from driving strong spiking responses.

An unusual and, we believe, critical feature of DL neurons is that they exhibit long-lasting spike threshold adaptation (*i.e*., threshold fatigue); our modeling suggests that this is due to Na^+^ channels exhibiting slow recovery from inactivation. The link between a sustained spike threshold increase and the slow inactivation of Na^+^ channels has previously been suggested for hippocampal CA1 pyramidal neurons (Henze and Buzsaki, 2001).

The standard ion channel repertoire of DL neurons appears at odds with the results of applying the Pozzorini protocol. This protocol resulted in complex and unpredictable spiking patterns including unexpected spike rate variations and, in some cells, resulted in long-term increases in cell excitability (Fig. 10). It is not clear whether these variable responses are due to the highly dynamic spike threshold adaptation kinetics across DL cells, or whether additional longer time scale processes are operating, e.g., very slow Ca^2+^-dependent changes in channel properties (Simms and Zamponi, 2014). Furthermore, DL is internally differentiated by gene expression (Harvey-Girard et al., 2007; Harvey-Girard et al., 2013; Ganz et al., 2014) and connectivity (Giassi et al., 2012c) which may also contribute to the highly variable responses. Developing a high quality model of DL cells is an essential next step in connecting the dynamics of the DL recurrent network (Trinh et al., 2016) to behavioral studies on spatial learning in the dark (Jun et al., 2016).

**Figure 10.**
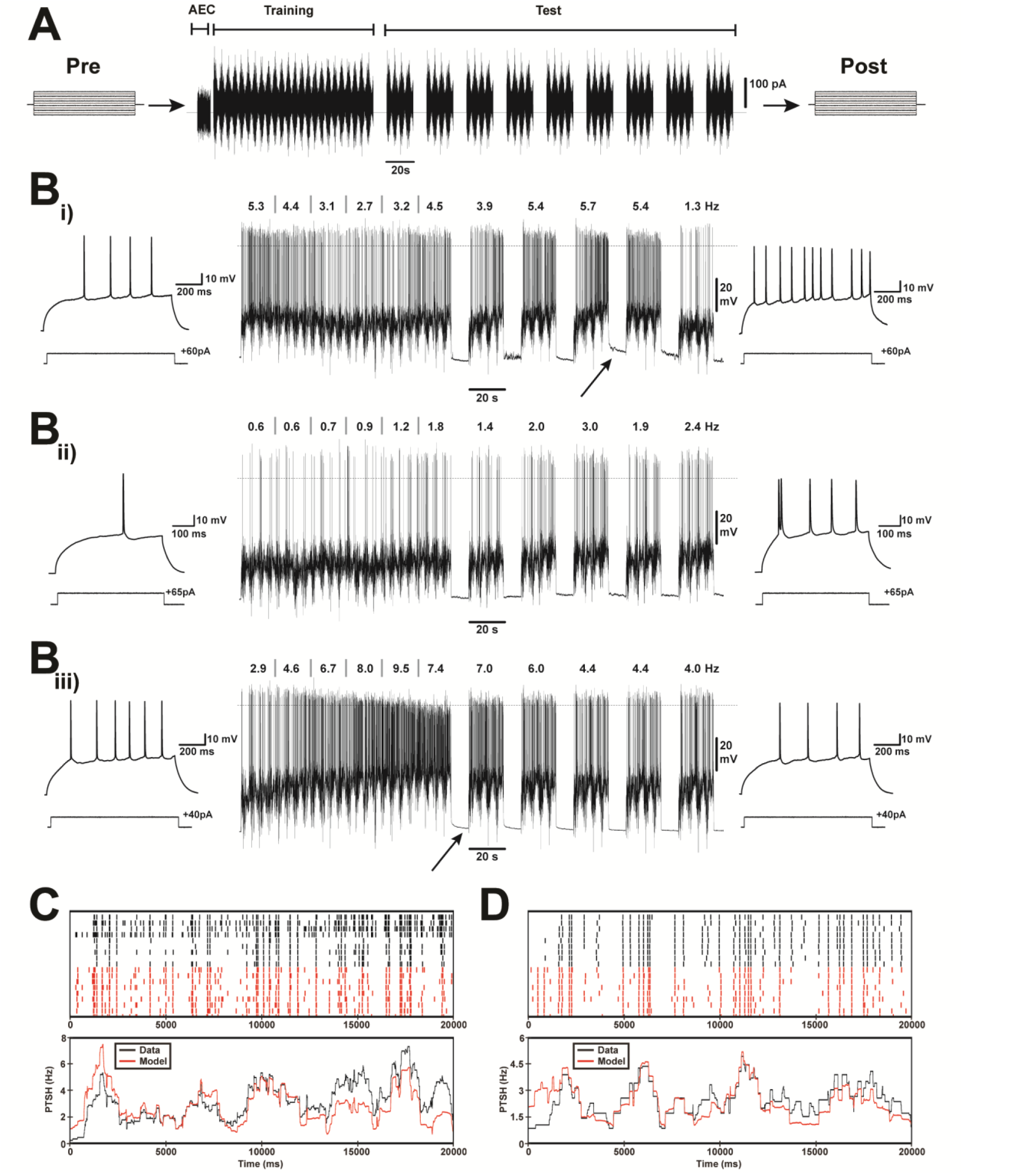
Fitting a Generalized Integrate and Fire (GIF) model to DL neuron spiking. **A.** General scheme of the parameter extraction protocol sequence (Pozzorini et al., 2015). A pre-test step current injection (500 ms) was first applied to assess the neuron’s firing rate response. Before the parameter extraction protocol, a 10 s subthreshold stimulus was applied to the cell in order to estimate the active electrode compensation filtering (AEC). This was then followed by the training phase (100 s) in which peri-threshold modulated noise was injected into the cell. After a 10 s pause, five test modulated noise stimuli were applied (20 s with interleaved 10 s pauses). The model parameters were extracted from the responses during the test phases (N = 13 cells). A post-test current pulse identical to the pre-test pulse was then applied to check on possible long-term changes evoked in the cell’s firing rate response. **B.** Example recordings from three different DL cells (*Apteronotus*, **i-iii**); the responses of these cells span the full range observed for all *Apteronotus* (N = 10) and goldfish (N = 3) cases. The dotted line near the peak of the spikes indicates a membrane potential of 0 mV. The numbers at the top of each trace represent the firing rate (in Hz) for successive 20 s intervals; 20 s was chosen to match the duration of the test stimuli. Only the first 5 test stimuli out of the total 9 stimuli are illustrated here since the spiking responses did not appreciably change after the 5^th^ test pulse. In all cases, the spiking rate is extremely variable during the training and test phases with periods of decreasing and increasing rates. The increases in firing rate were surprising given the increases in spike threshold evident in Fig. 8, which should make it harder for the cell to spike. The firing rate variability is not due to the deterioration of cell health since the spike height during the entire protocol does not change significantly. In some cases, during the near-threshold stimulus injection, the firing rate increases and this is accompanied by a change in membrane potential as highlighted by the black arrows (**i** and **iii**). In about half of the cases (**ii**; 6 out of 13 cells) the baseline membrane potential between tests remains constant. Furthermore, the noise stimulus would sometimes increase the excitability of the cell, as shown by an increase in firing rate during the post step-current injection compared to the pre step-current injection (**i, ii**; 5/9 cells) and in other cases, the excitability of the cell remained unchanged or decrease (**iii**; 4/9 cells). Overall, the response across different cells to the noise stimuli showed great variability; the response of each cell was variable as well, which presumably led to the inability of the GIF model to produce good consistent fits to the recorded data. **C.** Example good GIF model fit results for a cell displaying an increase in excitability after the Pozzorini protocol (model fit of the cell shown in 10B. **(i)**; Goodness of fit M* _d_ = 0.84. The boxed raster plot illustrates the spikes obtained during each of the nine noise test stimuli. The black dots represent the spikes obtained from the current-clamp experiments, while the red dots represent the predicted spikes obtained from the fitted GIF model. The lower box illustrates the peri-stimulus-time histogram (in Hz) of the model versus the one obtained from the recorded cell. **D.** Same as in C except this cell did not display any changes in excitability (Goodness of fit M* _d_ = 0.61).

### The biophysical properties of DL neurons suggest that they are sparse coders

The main properties that contribute to low DL neuron firing rates are the very depolarized spike threshold and hyperpolarized RMP (Table 2); these parameters are highly variable but typically lead to a large (∼32 mV) barrier that excitatory input must exceed to evoke spiking (Table 2). This contrasts sharply with the first order electrosensory pyramidal cells within the ELL. Their barrier from rest to spiking is a mere 4.9 mV (Table 2) and they can even respond to weak signals with discharge frequencies over 100 Hz. ELL pyramidal neurons also recover rapidly from spike induced increases in spike threshold, *i.e.*, threshold fatigue (10s of milliseconds, Chacron et al., 2007). We hypothesize that these properties are responsible for the ability of pyramidal cells to densely encode spatial and social electrosensory signals (Vonderschen and Chacron, 2011). The low barrier from RMP to spike threshold is also seen in primary auditory neurons and in layer 4 cells of the primary visual and somatosensory cortex (Table 2). Although no precise estimates are available, it appears likely that all these low level sensory neurons encode sensory input much more densely than neurons in the hippocampus.

**Table 2:** Difference in spike threshold and resting membrane multiple cell types.

|  | ‘hippocampus’ |  | L4 Sensory cortex |  | Primary Sens. cells |  |
| --- | --- | --- | --- | --- | --- | --- |
| Values in mV | DL cells | DG granule cells<sup>a</sup> | Barrel Field<sup>b</sup> | Visual Cortex<sup>c</sup> | ELL Pyr ON cells | DCN Pyr Cells<sup>f</sup> |
| Spike threshold | -44.2 | -40.8 | -45.1 | ~-63.5 | -62.9<sup>d</sup> | -48.1 |
| RMP | -76.2 | -74.7 | -63.0 | -72.0 | -67.8<sup>e</sup> | -62.7 |
| Threshold – RMP | 32.0 | 33.9 | 17.9 | 8.5 | 4.9 | 14.6 |

<sup>a</sup> Kowalski et al. (2016), *in vivo*, threshold: used point of 1<sup>st</sup> derivative which exceeded 20 V\*s<sup>-1</sup>.
<sup>b</sup> Yu et al. (2016), *in vivo*, threshold: used point of 1<sup>st</sup> derivative which exceeded 3 times the average 1<sup>st</sup> derivative.
<sup>c</sup> Wilent and Contreras (2005), *in vivo*, threshold: used peak of 2<sup>nd</sup> derivative, values were averaged from the data for preferred direction and non-preferred direction
<sup>d</sup> Mehaffey et al. (2008), *in vitro*, threshold: used point of 1<sup>st</sup> derivative which was 8 times greater than SD.
<sup>e</sup> Berman and Maler (1998a), *in vitro*.
<sup>f</sup> Li et al. (2013), *in vitro*, threshold: used point of 1<sup>st</sup> derivative which exceeded 10 V\*s<sup>-1</sup>.

Hippocampal neurons such as dentate gyrus granule cells are nearly silent at rest, and discharge very sparsely in response to the animal’s spatial location, *i.e.*, place field (Diamantaki et al., 2016). The low excitability in mature granule cells was shown to be partly due to the constitutive activity of GIRK channels (Gonzalez et al., 2018). We hypothesize that a similar mechanism is contributing to the low RMP of DL neurons in fish, which may partly explain why the difference between RMP and spike threshold is nearly identical in DL and DG cells (Table 2).

With the above examples in mind, we hypothesize that the key biophysical signatures of sparse coding are, for all neurons, a large gap between the RMP and the spike threshold in combination with spike threshold adaptation with long recovery timescales. We further hypothesize that DL neurons will sparsely encode the spatial relations required for memory guided navigation.

### Can the DL network transform PG sequential encounter time stamps to a spatial map?

Previous studies have investigated electrosensory spatial learning in a related gymnotiform fish (Jun et al., 2016; Fotowat et al., 2018). They showed that these fish can locate food relative to landmarks in the dark because, after learning, they rapidly navigated to the remembered food location during probe trials (no food). The electrosense is very local and, for most of their trajectory, the fish had no external sensory cues (Jun et al., 2016). This led Jun et al. to argue that, after leaving a landmark, the fish used path integration of speed and orientation signals to continuously update its current location and thus compute the trajectory to the remembered food location. Path integration information was assumed to potentially derive from lateral line receptors, vestibular afferents, proprioceptors and vestibular afferents. Bastian et al. (1995) has previously reported, in gymnotiform fish on brainstem proprioceptive neurons capable of signalling tail bending. Recently, Wallach et al. found PG neurons are responsive to continuous lateral line input, confirming a second potential source of information related to the fish’s speed. A recent study in the larval zebrafish has demonstrated that vestibular input can evoked strong and widespread activity in the telencephalon that, from the images presented, likely includes DL (Favre-Bulle et al., 2018). We now hypothesize that an encounter with a landmark triggers an autonomous ‘moving bump’ in the DL recurrent network and this is the primary driver for the fish’s estimation of its changing location during its landmark-to-food trajectories. While proprioceptive, lateral line and vestibular input are important, we now hypothesize they merely modulate the essential intrinsic DL network dynamics. We elaborate on this hypothesis below.

In gymnotiform fish, PG cells respond to object motion (electrosensory and visual; Wallach et al., 2018). Anatomical studies indicate that these responses are driven by tectal input (Giassi et al., 2011). The gymnotiform tectum maintains a topographic representation of electrosensory input and tracks continuous object motion (Bastian, 1982). PG neurons generate a major transformation of their tectal input - the majority of PG motion sensitive cells lose topographic information and respond over the fish’s entire body but only to object motion start (all cells) and stop (some cells) and not the intervening continuous motion (Wallach et al., 2018). Wallach et al. proposed that, during navigation in the dark, these PG cells will respond transiently when any part of the fish’s body first encounters a landmark (or food), *i.e.*, the response of the cell when the experimenter moves an object towards the fish is equivalent to its response when the fish moves near a landmark or food. Wallach et al. further proposed that the time interval between encounters could be ‘read out’ from the change in second versus first encounter firing rates of a subset of DL cells.

In the following discussion, we borrow extensively from work on ‘time’ and ‘place’ cells in the mammalian hippocampus using, in particular, the very thorough papers of Kraus et al. (2013) and Pastalova et al. (2008) as well as the related theoretical papers of Itskov et al. (2011) and Rajan et al. (2016). Kraus et al. describe hippocampal neurons that respond at specific times during a rat’s motion on a treadmill. These experiments carefully dissociated time from place so that the authors were able to demonstrate the existence of ‘time cells’, traditional place cells as well as cells with information on both the time and distance travelled. Pastalkova et al. and Itskov et al. had previously argued that sequential activation of cell assemblies is internally generated by hippocampal dynamics and can give rise to time cells independent of sensory input. Kraus et al. extended this hypothesis and argued that their time cells were driven by both internal network dynamics and external cues such as treadmill speed.

The theoretical papers of Itskov et al. and Rajan et al. asked: how might the intrinsic activity of a neural network result in the sequential activation of neuron assemblies, e.g., time cells? Both papers started with the same core architecture – a local excitatory recurrent network that, once activated, was capable of sustained discharge. This is the ‘bump attractor’ hypothesis originally formulated to explain the sustained activity of neurons during a working memory task (Wang, 1999; Wimmer et al., 2014). The theoretical analysis of Wang (1999) demonstrated that slow excitatory synapses, *i.e.*, mediated by NMDA receptors, were required for bump dynamics. Both Itskov et al. and Rajan et al. generated ‘cell assembly sequences’ by destabilizing the bump attractor dynamics. Itskov et al. accomplished this by introducing spike threshold adaptation with a long recovery time constant. In contrast, Rajan et al. destabilized the bump by introducing asymmetries in synaptic strengths within the attractor so that the attractor dynamics would generate a sequential activation of the cell assembly; a process which necessitated both recurrent connections and external input. In both cases, sequential activation of neurons within the cell assembly are able to produce time cells or other sequential outputs. A recent paper (Heys and Dombeck, 2018) has also suggested that ‘time cells’ of the entorhinal cortex might be generated by moving bumps in entorhinal recurrent attractor network (Zutshi et al., 2018). This paper did not, however, explicitly discuss the mechanism by which the putative ‘bumps’ would move.

Our earlier work (Trinh et al., 2016) demonstrated that DL contains excitatory local recurrent networks; our earlier work had already demonstrated that DL is highly enriched in NMDA receptors (Harvey-Girard et al., 2007). Trinh et al. therefore hypothesized that the DL recurrent network supported bump attractor dynamics capable of memory storage. Our ‘noisy’ cells suggest that the recurrent connections within DL are, in fact, capable of supporting autonomous discharge. We have now demonstrated that DL neurons exhibit the same threshold adaptation utilized in the Itskov et al. model thus suggesting that the putative DL bumps may not be stable attractors. We have not yet studied the properties of either PG-derived or intrinsic synapses in DL and therefore cannot evaluate whether Rajan et al.’s architecture might apply. In accordance with the Itskov model, we hypothesize that DL contains unstable bump attractor neural networks that are capable of supporting autonomous sequential activation and thus DL time cells. We assume that, when the fish initially encounters a landmark, the resulting electrosensory-evoked transient discharge in a subset of PG neurons triggers activity in a small region of DL (Giassi et al., 2012c). This activity will then propagate through a subset of the DL network forming a cell assembly temporal sequence (time cells). Following Kraus et al., we further hypothesize that the sequential activity in this network is modified by ongoing self-motion sensory input – the vestibular, lateral line and proprioceptive input mentioned above. These inputs provide the path integration signals that converts the time cell sequence to a location cell sequence. In functional terms, we propose that the propagation of neural activity in the DL network represents the fish’s estimate of where it is located along the trajectory between a landmark and food. When the fish reaches the food (or another landmark), PG neurons would again discharge to signal the total time/distance travelled (Wallach et al., 2018) and the potential start of a new trajectory. In this model, learning a trajectory from a particular landmark to food would consist of strengthening the synaptic connections of the ‘moving bump’ induced by that landmark so as to represent the time/location sequence leading from the landmark to food. Such strengthening might result in Rajan et al. type mechanism in which directed bump movement was now also a consequence of asymmetric synaptic strengthening.

Our hypotheses are at the moment not testable, because testing would require population recording from or visualizing activity across a large portion of the DL network. What is needed is a teleost that is transparent when adult, whose neurons express a genetically encoded calcium indicator (e.g., gCamp6) and whose pallium might be activated by ethologically relevant transient signals. Fortunately, such a model system has recently become available (Schulze et al., 2018) and may permit direct tests of our hypotheses.

## Materials and methods

For the following experiments, we used two closely related *Apteronotid* fish (*A. leptorhynchus* and *A. albifrons*), a suborder of the gymnotiform family, as well as *Carassius auratus* (goldfish). The brains of *A. leptorhynchus* and *A. albifrons* cannot be readily distinguished; these species have been used interchangeably in previous anatomical studies (e.g., Carr et al., 1982) and the processing of electrosensory input appears to be nearly identical in these species (Martinez et al., 2016). Goldfish were included in this study for three reasons: First, we found that *Apteronotus* DL cells were challenging to maintain in slice preparation, whereas goldfish DL cells were more robust, yielding higher success rates on our lengthier protocols involving pharmacological manipulations. Second, we wanted to check how our results generalized to non-electrosensory teleosts, given the very general mechanisms of sparse neural coding proposed in this article. Last, the critical behavioral experiments on the essential role of DL in spatial memory were done in goldfish (Rodriguez et al., 2002), setting a precedent in the literature; further, the first *in vivo* DL recordings have also been carried out in goldfish (Vinepinsky et al., 2018). As demonstrated in the results, our conclusions apply equally well to each of these species and are therefore directly relevant to spatial learning across a broad range of teleost fish.

Prior to use, the *Apteronotus* fish were kept in heated aquariums at 28°C, while goldfish were kept in aquariums at 22°C (room temperature). All procedures were approved by the University of Ottawa Animal Care Committee and follow the guidelines issued by the Society for Neuroscience.

### Slice preparation

Prior to the dissection, adult male and female fishes were anesthetized in oxygenated water containing 0.2% 3-aminobenzoic ethyl ester (tricaine methanesulfonate, Aqua Life, Syndel Laboratories, Canada). As the skull was being removed, ice cold oxygenated (95% O_2_, 5% CO_2_) artificial cerebrospinal fluid (ACSF: 130 mM NaCl, 24 mM NaHCO_3_, 10 mM Glucose, 2.5 mM KCl, 1.75 mM KH_2_HPO_4_, 1.5 mM CaCl_2_, 1.5 mM MgSO_4_, 295 mOsm, pH 7.4), containing 1 mM of kynurenic acid (Millipore Sigma, Canada), was dripped onto the fish’s brain. The brain was then carefully removed and submerged in a petri dish containing ice-cold ACSF with kynurenic acid. Once the brain was removed, it was placed in an ice-cold cubic mold, to which oxygenated ACSF mixed with 2.5% low-melting agarose (Millipore Sigma, Canada) was added. After the agarose has polymerized, an initial cut was performed in order to separate the telencephalon from the rest of the brain. Subsequently, 300 µm thick coronal brain slices of the telencephalon were obtained using a vibratome, as previously described (Trinh et al., 2016). For goldfish dissections, the thick optic nerves underneath the brain had to be severed with micro scissors before the brain was removed and placed in a petri dish containing ice-cold ACSF. The rest of the dissection was done in the same manner as in *Apteronotus* (see Trinh et al., 2016). Brain slices containing the dorsal lateral telencephalon (DL) were then transferred into a continuously oxygenated slice incubation chamber containing ACSF where they were left to rest for 30-60 minutes.

### In vitro recordings

After the incubation period, brain slices containing DL were transferred to the recording chamber where oxygenated ACSF was constantly perfused at a flow rate of 3 mL/min. Recordings were performed at room temperature (23-24°C). We used fire-polished borosilicate glass micropipettes (Sutter Instruments, CA) with resistances ranging between 8-14 MΩ. The intracellular solution contained (in mM): 130 K-Gluconate, 10 KCl, 10 HEPES, 4 NaCl, 4 Mg-ATP, 10 phosphocreatine, 0.3 Na-GTP, with an osmolality of 295 mOsm, and a pH of 7.2 for weakly electric fish recordings. A silver wire plated with silver chloride was used as a ground. For goldfish experiments, slightly different extracellular and intracellular solutions were used (in mM): ACSF - 108 NaCl, 24 NaHCO_3_, 10 Glucose, 2.5 KCl, 1.25 KH_2_HPO_4_, 1.5 CaCl_2_, 1.5 MgSO_4_, 2 HEPES, 260 mOsm (adapted from Palmer, 2006). Intracellular solution - 110 K-Gluconate, 10 KCl, 18 HEPES, 4 Mg-ATP, 10 phosphocreatine, 0.3 Na-GTP, 265mOsm, pH 7.2 (adapted from Palmer, 2006). To visualize the neurons, slices were imaged under differential interference contrast (DIC) optics using a CMOS infrared camera (Scientifica, UK) directly connected to the rig computer. The recording signals were amplified using a Multiclamp 700B (Axon Instruments, CA), while the signal was filtered at 3 kHz and digitized using a Digidata 1550 (Molecular devices, CA). The whole-cell recording data was acquired using the PClamp 10.6 software (Molecular devices, CA). All recordings were performed in current-clamp mode. Only cells that required a minimal holding current less than - 50 pA were included in the study, allowing to stabilize the cell near the average resting membrane potential (approximately −75 mV, see Fig. 2E). The maximal recording time after the dissection was 4-5 hours. Once the whole-cell configuration was obtained, the resting membrane potential was recorded for 10 seconds and the cells were injected with current steps, which typically range from 500 to 1000 ms and from −60 to +60 pA, except where otherwise noted. For our ramp current protocol, we injected two different ramp currents at different inter-stimulus time intervals ranging from 50 to 1000 ms. Although both ramp stimuli have the same slope, the first ramp current was always two-fold stronger than the second ramp since the first ramp current had to evoke multiple action potentials while the second one only had to evoke one action potential. As such, the magnitude of the second current injection had to be adjusted for each cell since the rheobase for each cell is different and the magnitude of the first ramp was then adjusted according to the second ramp. Healthy cells were usually held for 30 to 60 minutes.

### Pharmacology

To test for the presence of fast and persistent sodium channels in DL neurons (Berman et al., 2001), we first patched the cell and injected a standard 500 ms current step before applying 20 µM TTX (Tetrodotoxin; Abcam, MA) locally near the recording site by pressure injection. To further investigate the presence of a persistent sodium channel, we also applied 5 mM QX-314 (Lidocaine N-ethyl bromide; Millipore Sigma, Canada) via the intracellular recording solution in order to block sodium (Salazar et al., 1996) and other channels (e.g., certain K^+^ channels and Ca^2+^ channels, see Results) (Alreja and Aghajanian, 1994; Perkins and Wong, 1995; Talbot and Sayer, 1996).

Calcium-activated potassium channels SK1/2 are both expressed in DL (Ellis et al., 2008). We used our standard current step protocol to evoke spikes in patched DL cells and bath applied an SK channel blocker 30 µM UCL (6,12,19,20,25,26-Hexahydro-5,27:13,18:21,24-trietheno-11,7-metheno-7*H*-dibenzo [*b,n*] [1,5,12,16] tetraazacyclotricosine-5,13-diium dibromide; Tocris, Bio-Techne, Canada) (Harvey-Girard and Maler, 2013). We also locally applied 1 mM EBIO (1-Ethyl-2-benzimidazolinone; Abcam, MA), a SK channel agonist near the brain slice by pressure injection (Ellis et al., 2007). Finally, we patched neurons using a slightly altered internal solution that contained 10 mM BAPTA (Millipore Sigma, Canada) in order to chelate intracellular calcium.

### RT-PCR

G protein-coupled inwardly-rectifying potassium channels (GIRK) 1-4 mRNA sequences were identified from *A. leptorhynchus* brain transcriptome data (Salisbury et al., 2015). Two degenerate PCR primers were designed to bind all GIRK isoform sequences (Forward: CTGGTGGACCTSAAGTGGMG; reverse: TTCTTGGGCTGNGNAGATCTT). Five *A. leptorhynchus* fish were anesthetized with tricaine methanesulfonate (Aqua Life, Syndel Laboratories, Canada) and then sacrificed by cervical dislocation while being fed oxygenated water containing the anesthetic. Different regions of the brain (DL, tectum/torus, subpallium, cerebellum, ELL, hindbrain) were dissected in ice-cold ACSF, collected and preserved on dry ice. All tissues were weighed, and homogenized in Trizol to purify total RNA (Millipore Sigma, Canada). First-strand cDNAs were then generated by using the RevertAid H Minus First Strand cDNA Synthesis Kit (Fermentas, MA). Degenerate polymerase chain reaction (PCR) was performed using the DreamTaq, according to the manufacturer recommendations (Thermo Scientific, MA), with the primers mentioned above. On an agarose gel, the amplicon expected bands were 344bp.

### Data analysis

All the recording data was first visualized in Clampfit (Molecular Devices, MA) before being transferred into Matlab (Mathworks, MA) for subsequent analysis with custom scripts. To reduce the likelihood of analysing unhealthy cell responses, only cells which produced spikes that cross a data-driven threshold of −5mV were included in the analysis. Cells that showed significant membrane noise, *i.e*., a variance greater than 0.5 mV^2^, were used to construct Fig. 2G but were excluded from any additional analysis. For the analysis of the resting membrane potential (Figure 1), only cells that did not require a holding current to stabilize were included in this analysis. The membrane time constant was measured by fitting an exponential function to the neuron’s recovery to equilibrium following injection of a negative step current. The spike amplitude was measured by two methods: first, as the difference between the spike height and the spike threshold and, second, from the difference between the spike height and the resting membrane potential. In order to estimate the spike threshold, we used the method of Azouz et al (2000) which defined the spike threshold as the voltage corresponding to an empirically defined fraction (0.033) of the peak of the first derivative. This first derivative method was later shown to be slightly better than the second derivative method (Sekerli et al., 2004) previously used for hindbrain electrosensory neurons (Chacron et al., 2007). The threshold for the broad Ca^2+^ spikes were determined visually in Clampfit since the rate of change of the Ca^2+^ spike was too slow to be visualize with either the 1^st^ or 2^nd^ derivative of the membrane potential. The spike width was calculated by measuring the half-width at half-maximum. The voltage, as a function of injected current (I-V curves), was obtained in Clampfit using sub-threshold traces and averaged in order to reduce the variability across cells caused by the holding current. The input resistance was obtained by calculating the average slope of the I-V curve across all cells. The after-hyperpolarization potential (AHP) amplitude was measured as the difference between the spike threshold and the minimum value of the AHPs. If the recording trace contained a burst or spike doublet, then the AHP would be measured on the following spike, since a doublet would typically induce an especially large AHP. The cell’s average firing rate was calculated as the number of spikes divided by the duration of the stimulus. The delta spike height was calculated as the difference in spike height between the n^th^ spike and the first spike of an evoked spike train. The inter-spike interval (ISI) was measured as the time between the first two spikes of the spike train induced by a current step injection, while the delta time was calculated as the difference between the time of the first AHP and the time of the n^th^ AHP. The delta AHP was obtained by subtracting the first spike’s AHP amplitude from the second spike’s AHP amplitude. The delta threshold was obtained in a similar fashion. All error bars were determined using the standard error of the mean. Wherever applicable, the statistical significance was determined using either one-way ANOVA, two-way ANOVA or the paired t-test, where p < 0.05 is considered significant.

### Inactivating exponential integrate and fire model (iEIF)

In order to illustrate the putative role of slow sodium channel inactivation on the observed and variable spike threshold in DL cells, we sought a minimal neuron model that incorporates an abstraction of sodium channel dynamics. The inactivating exponential integrate and fire neuron (equation [1]; Platkiewicz and Brette, 2011) provides a distilled representation of sodium channel activation via an exponential amplification of the membrane voltage (*V*), which is attenuated by fast and slow inactivation variables (*h*_*f*_ and *h*_*s*_). These sodium channel inactivation terms further affect the dynamic threshold for spike generation, *θ*, whose initial value *V*_*T*_ reflects no inactivation at the resting membrane potential (equation [2] and [3]; *h*_*f*_ = *h*_*s*_ = 1) (Platkiewicz and Brette, 2010). Although the exponential approximation does not realistically capture the full action potential waveform, which spans a large voltage range, it is valid for voltages near spike initiation. Importantly, this approximation permits the differential equation for the variable spike threshold, *θ*, to be simply expressed by sodium channel properties described in equations [2] and [3] (Platkiewicz and Brette, 2010, 2011).

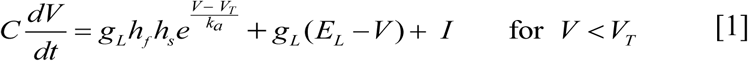

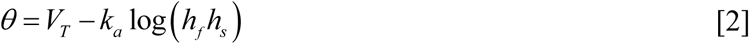

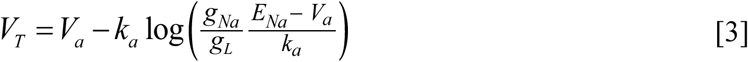

As in the work of Platkiewicz and Brette (2011), the membrane time constant, *τ* = *C* / *g*_*L*_ = 5 ms, was introduced for our simulations. Given that the specific membrane capacitance is about 0.9 µF/cm^2^ for practically all neuron types (Gentet et al., 2000), the leak conductance is constrained to be *g*_*L*_ = 0.18 mS/cm^2^ and the input current, *I* = 3.8 nA, is scaled by the associated membrane resistance (5.56 MΩ). The leak current reversal potential was set to *E*_*L*_ = −55 (Platkiewicz and Brette, 2011). When the membrane voltage reaches *θ* at time *t*, a spike is generated and *V* (*t*^+^) is reset to the resting membrane potential, *V*_*r*_ = −70 mV. The average threshold for the first spike in DL neurons was −42.96 ± 0.5 mV (N = 42 spikes). To obtain an approximate match between *V*_*T*_ and this value, we kept the sodium activation slope, *k*_*a*_ = 4 mV, and reversal potential, *V*_*a*_ = −38.6 mV, at the empirically justified values used by Platkiewicz and Brette (2011). We then set the sodium conductance to *g*_*Na*_ = 0.036 mS/cm^2^ to achieve a value of *g*_*Na*_/*g*_*L*_ = 0.2, near the range of Platkiewicz and Brette (2011). We assume this slightly lower value in our model is a reflection of low sodium channel density. Consistent with this assumption, DL neuron axons are very thin and possibly unmyelinated (Trinh et al., 2016), which presumably explains the high DL neuron threshold. The sodium channel reversal potential was kept at a standard *E*_*Na*_ = 50 mV. When substituted into equation [3], the above parameter set yielded an initial threshold of *V*_*T*_ = −44.6 mV (Fig. 8E; ii) and gave particularly close agreement with the *Apteronotus* data (−44.5 ± 0.2 mV; Fig. 4E).

Drawing on the Hodgkin-Huxley formalism, the inactivation variables, *h*_*f*_ and *h*_*s*_, evolve according to equations [4-5], where *h*_∞_ is a Boltzmann equation with inactivation parameters *V*_*i*_ = −63 mV and *k*_*i*_ = 6 mV [6]:

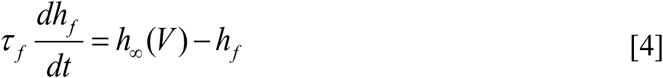

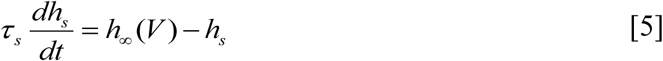

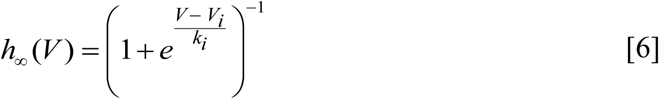

The parameters *τ* _*f*_ (fast inactivation timescale) and *τ* _*s*_ (slow inactivation timescale) are of particular interest to the model and to our results. To determine *τ* _*f*_, the average time between a short burst of two DL spikes (doublet) was measured at the beginning of the recorded voltage trace, yielding 15.38 ± 0.6 ms (N = 144 doublets). Selecting *τ* _*f*_ = 15 ms, we note that the model generates spikes at a frequency of 64.7 ± 7.8 Hz, consistent with the data mean. We assumed that a slow timescale of inactivation would lead to an increase of spike threshold with a correspondingly long timescale for recovery (see Discussion). To select *τ* _*s*_, we therefore noted that the threshold for DL cell spiking remains significantly increased for at least 300 ms when stimulated; therefore, *τ* _*s*_ is likely on the order of 10^2^ milliseconds. A more direct estimate gave a mean decay time constant (τ_exp_) of ∼640 ms for the slow recovery (see ramp protocol results below). Therefore, we selected *τ* _*s*_ = 500 ms, which is a conservative value, given slow inactivation is typically >1 s and longer timescales would only further strengthen our hypotheses (Itskov et al., 2011).

Note that we omitted Ca^2+^ currents and the resulting SK channel mediated AHP since its duration is less than the typical interspike interval of DL neurons. When simulating the model, subthreshold Gaussian noise, *N* (0,1), was added to equation [1] and scaled by a factor *σ* = 0.5. The stochastic forward Euler method was used as the numerical solver. Code will be made available upon request to the corresponding author.

### Fitting the DL neuron responses to a generalized-integrate and fire (GIF) model

We used a protocol developed for cortical pyramidal cells (Pozzorini et al., 2015) to model spike generation in DL cells. Spike time extraction was done by injecting a modulated noise stimulus in three distinct phases. In the first phase, 10 seconds of sub-threshold noise was injected into the cell in order to estimate the active electrode compensation (AEC), which corrects for the bias in voltage drop caused by the pipette. This phase is followed by a 2 s delay where no current is injected, after which the cell is subjected to 120 seconds of near-threshold noise stimulus, which was generated according to an Ornstein-Uhlenbeck process whose variance is modulated by a 0.2 Hz sinusoid. All the parameters used to generate this noise stimulus were taken from Pozzorini et al. (2015) except for the intensity of the input which had to be reduced 4-fold, presumably because of the far smaller size of DL cells compared to mammalian cortical pyramidal cells (Giassi et al., 2012b; Trinh et al., 2016). This training phase was then followed by the test phase, over which the cell received 9 different realizations of the same near-threshold oscillating noise stimulus for 20 s intervals, with 10 s of rest in between. The purpose of the training phase was to extract parameters that were then used to construct a GIF model of the DL neuron. The test trials were carried out in order to compare the evoked spiking pattern with that generated by the GIF model (Fig. 7A). The entire methodology of this section was adapted from the methods used in Pozzorini et al. (2015) and the Python code used for this section was obtained from the authors’ GitHub page, however, all simulations were done in Matlab. The firing rate of these cells was then determined by counting the number of spikes within 20 s bins throughout the training protocol; this interval was chosen to match the duration of test phases. Before applying this fitting protocol, we tested the patched cell’s health by injecting either 500 ms or 1000 ms current steps, which is defined as the “pre” period. This same current step was then reapplied during the “post” period, after the cell underwent the Pozzorini et al. (2015) fitting protocol, in order to confirm that the cell’s health had not deteriorated.

All panel figures were initially compiled in OriginPro 9.0 (OriginLabs, MA) and the final figures were assembled in Adobe Illustrator (Adobe Systems, CA).

## Acknowledgements

We would like to thank William Ellis for technical support and Maria Lambadaris for her help with the electrophysiological recordings. We also thank Jean-Claude Béïque and Timal Kannangara for their helpful discussions and suggestions. This research was supported by the CIHR 153143 and NSERC 211418 grant, both assigned to LM.

## Competing interests

The authors declare no competing financial or non-financial interests.

---

**Figure S1.**
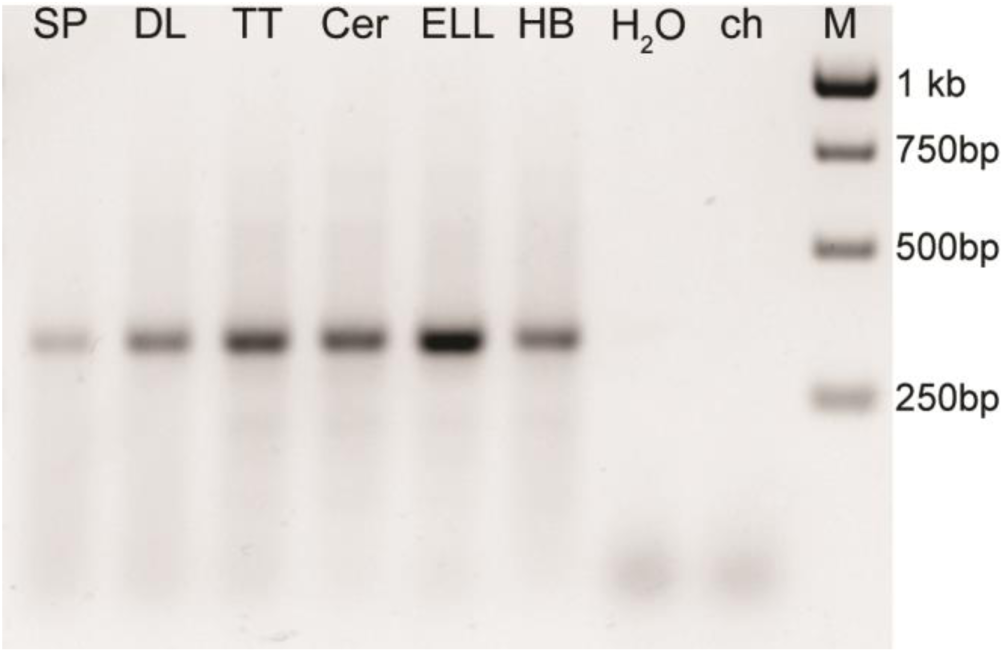
GIRK channel mRNA expression obtained from RT-PCR in the *Apteronotus* brain using pan-PCR primer pairs in conserved regions. GIRK channels are ubiquitously expressed albeit at variable levels. In particular they are expressed in DL. **Abbreviations**
SP: subpallium
DL: dorsolateral pallium
TT: Tectum/Torus
Cer: cerebellum
ELL: electrosensory lobe
HB: hindbrain
ch: chicken (negative control)
M: molecular marker

**Figure S2.**
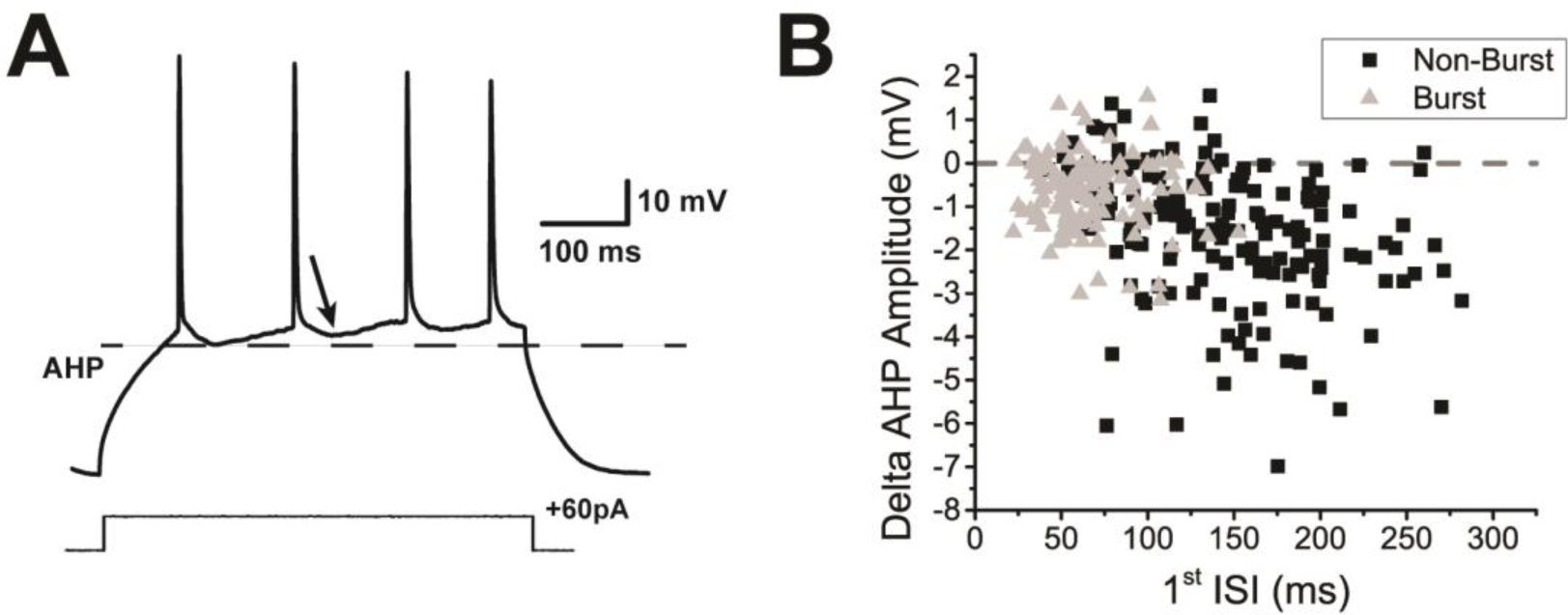
Current-evoked spiking decreases the AHP amplitude of DL neuron. **A.** Example trace of a DL neuron’s response to a +60 pA current step injection lasting 500 ms. Consecutive spiking causes the AHP amplitude to decrease when compared to the first AHP as emphasized by the arrow. A black dashed line is placed to coincide with the minimum of the first spike’s AHP. We hypothesize that the decrease in AHP amplitude is due to a reduction in Ca^2+^ influx (*i.e.*, Ca^2+^-dependent Ca^2+^ channel inactivation) and subsequent reduction in SK channel opening. **B**. The decrease in AHP amplitude between the second and first spikes is plotted as a function of the time interval between the first two spikes similarly to Fig. 8B. Each black square represents a spike pair taken from a trace which did not contain a burst (total of 160 non-burst spike pairs), while each grey triangle represents a spike pair taken from a trace which contained a burst at the beginning of the trace (total of 117 burst spike pairs). The majority of the AHPs are reduced throughout the 300 ms test period without any evident recovery trend.

**Figure S3.**
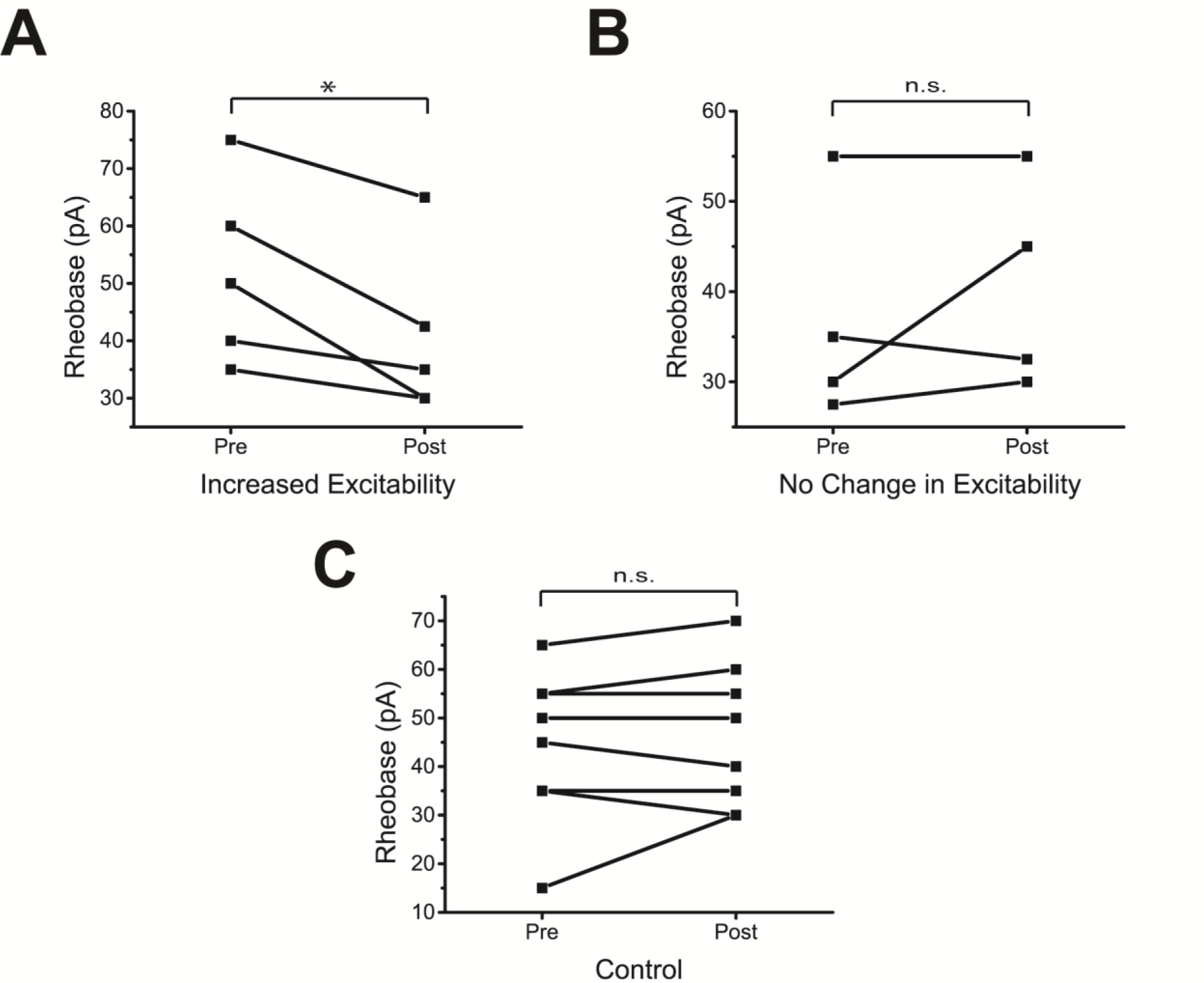
Changes in rheobase current before and after the application of the Pozzorini protocol **A.** DL neurons that displayed an increase in firing rate also showed a decrease in rheobase current after the Pozzorini protocol. **B**. Same as in A except that cells showing no change in excitability also did not display a significant change in rheobase current after the Pozzorini protocol. **C**. As a control, the rheobase was measured before and after a wait time that is equivalent to the duration of the Pozzorini protocol (7 mins). Following the wait time, the rheobase did not significantly change.

